# Deciphering the Genome Protection Roles of Autophagy in Primary Human Dermal Fibroblasts (HDFs) against Ultraviolet-(B) –Induced Skin Photodamage

**DOI:** 10.1101/2020.09.28.316273

**Authors:** Sheikh Ahmad Umar, Sheikh Abdullah Tasduq

**Affiliations:** Academy of Scientific and Innovative Research (AcSIR), Ghaziabad, 201002, India, Jammu Campus; Pharmacokinetics-Pharmacodynamics (PK-PD) and Toxicology Division, CSIR-Indian Institute of Integrative Medicine, Jammu Tawi, 180001, Jammu and Kashmir, India

**Author notes:** **Correspondence:** Sheikh Tasduq Abdullah ***PK-PD and Toxicology Division**, CSIR-Indian Institute of Integrative Medicine (IIIM), Council of Scientific and Industrial Research (CSIR), Canal Road Jammu Tawi, Jammu and Kashmir India., Ext-332.

**Keywords:** Autophagy, Ultraviolet Radiation, Primary Human Dermal Fibroblasts, Genotoxicity, Oxidative Stress, Endoplasmic Reticulum Stress, ATG7

## Abstract

Ultraviolet-B (UV-B) exposure to skin causes photo-damage and acts as the primary etiological agent in photo-carcinogenesis. UV-B exposure induces photodamage in epidermal cells and is the major factor that challenges skin homeostasis. Autophagy allows fundamental adaptation of cells to metabolic needs and stresses. Cellular dysfunction is observed in aged tissues and in toxic insults to cells that undergo through stress. Conversely, promising anti-aging strategies aimed at inhibiting the mTOR pathway has been found to significantly improve the aging related disorders. Recently, autophagy has been found to positively regulate skin homeostasis by enhancing DNA damage recognition. Here we investigated the Geno-protective roles of autophagy in UV-B exposed primary HDFs. We found that improving autophagy levels in HDFs regulates UV-B mediated cellular stress by decreasing the formation of DNA photo adducts, alleviates oxidative and ER stress response and by regulating the expression levels of cell cycle regulatory proteins P21 and P27. Autophagy also prevents HDFs from UV-B -induced nuclear damage as is evident from Tunnel assay and Acridine Orange/Ethidium Bromide co-staining. Salubrinal, (an eIf_2_α inhibitor) significantly decreases the DNA damage response in HDFs. P62 silenced HDFs show enhanced DNA damage response and disturbs the tumor suppressor axis PTEN/pAKT towards damage whereas ATG7 silenced HDFs reveal an unexpected consequence by decreasing the UV-B -induced DNA damage compared to UV-B treated HDFs. Together, our results suggest that autophagy is essential in protecting skin cells from UV-B radiation -induced photo-damage and holds great promise in devising it as a suitable therapeutic strategy against skin photo-damage.

**Highlights:**

1. Autophagy is an immediate molecular event induced following exposure of primary HDFs to UV-B –irradiation
2. Autophagy offers pro-survival capacity to HDFs under UV-B induced genotoxic stress
3. Autophagy regulates DNA Damage Response via regulation of oxidative and ER stress in UV-B exposed HDFs
4. Relieving ER stress response offers significant protection to primary HDFs from UV-B by decreasing the DNA damage
5. Autophagy deprivation to HDFs via P62 silencing potentiates UV-B -induced DNA damage response
6. ATG7 silencing in UV-B exposed HDFs unexpectedly alleviates the DNA Damage Response in primary HDFs

## 1. Introduction

Skin being the external covering to body protects internal organs from outside environmental insults including that from the adverse effects of ultraviolet (UV) irradiation [1]. Though ubiquitous presence of solar radiation is essential for survival to different life forms on earth but excessive exposures, either, acute or chronic, leads to skin photoaging, oxidative damage and malignancies constituting photodamage and photo-carcinogenesis through its UV-B portion of spectrum [2]. Macroautophagy (hereafter referred to as autophagy) at basal levels protect cells from stress and nutrient deprivation during starvation conditions and thereby maintains tissue homeostasis [3]. Cellular autophagy levels can be improved chemically especially in skin cells to maintain tissue homeostasis in response to diverse range of physiological and pathological stresses, including from solar UV-B -irradiation [4–6]. Dysfunctional autophagy has been associated with multiple human diseases, such as metabolic diseases, cardiovascular diseases, aging, neuro-degeneration, infectious diseases and in cancer and attempts are being made to use autophagy as selective therapeutic intervention in different disease conditions based on its differential roles it performs to maintain tissue homeostasis [7, 8]. Role of autophagy is context dependent and can be both oncogenic and tumor suppressive [9] either promoting or suppressing tumorigenesis through selective degradation of SQSTM1/p62, an autophagy signalling and adaptor receptor that promotes inflammation, cell proliferation and migration [10, 11], through removal of cellular debris to prevent genomic damage, or by promoting DNA repair in response to ionizing radiation-induced DNA double-strand breaks in mammalian cells to restore tissue homeostasis [11, 12]. In either way, the role of autophagy is to protect cells from external insults that disturb the cellular homeostasis. Recently, it has been found that autophagy regulates nucleotide excision repair (NER) that eliminates DNA base lesions induced by solar UV-B radiation, including cyclobutane pyrimidine dimers (CPD) and pyrimidine-(6-4)-pyrimidone photoproducts (6-4 PP) [13–15], in an attempt to restore tissue homeostasis and prevent tumor formation in cells. It has also been found that recruitment of DDB2 to UV-induced CPD sites is significantly impaired in autophagy deficient cells. In mice, rapamycin, (pharmacological autophagy inducer) was found to decrease the UVB-induced tumorigenesis while the inhibitor Spautin-1 increases it [16, 17]. These findings cite the critical role of autophagy in maintaining proper NER activity and suggest a new tumor-suppressive mechanism of autophagy in tumor initiation and regulation. Previously, we have reported from our own lab that UV-B induced Ca^2+^ deficit within ER lumen is mediated by immediate oxidative stress induced upon UV-B – irradiation to skin cells followed by subsequent photo-damage response. Insufficient Ca^2+^ reserves within ER lumen develop ER stress leading to Unfolded Protein Response (UPR) in skin cells that ultimately disturb the internal cellular homeostasis [18]. We have also reported in another of our study that natural anti-oxidant agent Glycyrrhizic acid (GA) potentially modulates cellular response in UV-B –irradiation to primary HDFs notably due to the reduction of oxidative stress-induced DNA damage, modulation of inflammation and regulation of pro-apoptotic and autophagic signaling possibly by adjusting the adaptation of cells to stresses [19]. Despite these preliminary findings, the role of autophagy in UV-B induced photo-damage response in unclear and requires further studies to unearth the facts. In line with these findings, here we attempted to investigate the roles of autophagy in regulating tissue homeostasis notably under genotoxic stress on UV-B radiation exposure to HDFs. We found that autophagy levels are impaired at higher intensity of UV-B -irradiation 30 mJ/cm^2^ with physiological exposure of environmental UV-B – irradiation being 10 mJ/cm^2^. Enhancing autophagy with pharmacological activator Rapamycin significantly reduces the formation of oxidatively induced DNA photo adducts CPD’s and 6,4, PP’s, restores the ER calcium levels and alleviates oxidative ROS species in HDFs in 6h UV-B post –irradiation exposure. We found that autophagy is an immediate molecular event following UV-B exposure to HDFs and is induced within 1-6h UV-B post-irradiation interval and with increasing the time interval the autophagic flux vanishes. Further, relieving ER stress response with Salubrinal prevents oxidative DNA damage in UV-B exposed HDFs. It was also revealed through florescent microscopic studies that Rapamycin treatment significantly alleviates Tunnel positive cells indicating nuclear damage and EtBr positive cells indicating apoptosis in UV-B exposed HDFs. The GFP/RFP LC3B puncta assay depicts greater puncta positive cells in Rapamycin treated cells than in UV-B treated alone showing enhanced autophagy flux in rapamycin treated cells upon UV-B -irradiation to HDFs. Rapamycin treatment significantly decreases the expression profile of key cell cycle regulators P21, P27 and DNA damage response pathway proteins DDB2, pP53 indicating that autophagy has cell protective roles in UV-B induced photo-damage. We also found least pGH_2_AX foci in Rapamycin treated cells than in UV-B exposed which is the main DNA damage sensor and among the first proteins that are recruited at the sites of DNA damage. P62 silencing confirmed our findings as the pGH_2_AX foci were significantly increased in P62 silenced cells than in UV-B exposed only in confocal microscopy, indicating that autophagy deprivation to UV-B exposed HDFs increases the damage response whereas increasing the flux decreases the DNA damage response positively. We further found that the tumor suppressive axis PTEN/pAKT having roles in preventing tumorigenesis is also significantly disturbed in P62 silenced cells. ATG7 silencing reveal an unexpected consequence by decreasing the DNA damage in UV-B exposed HDFs indicating that the roles of autophagy in protecting skin cells against radiation induced genotoxic stress are diverse and through differential protein expression of autophagy related genes. Together, our results suggest that improving autophagic flux in HDFs significantly alleviates the UV-B mediated DNA damage response and holds great promise in devising it as a suitable therapeutic and cosmeceutical strategy in combating radiation induced skin photo-damage disorders.

## 2. Materials and Methods

### 2.1 Chemicals

Human primary Dermal Fibroblast cell line from the juvenile foreskin (HDF) and primary fibroblast expansion media was obtained from HiMedia, Mumbai India. Fetal bovine serum (FBS), penicillin–streptomycin, trypsin–EDTA, 3-(4, 5-dimetylthiazol-yl)-diphenyl tetrazolium bromide (MTT), phosphatase-protease cocktail, RIPA buffer, H_2_DCFDA dye were purchased from Sigma–Aldrich Chemicals (St. Louis, MO). Antibodies against P62, BECN1, ATG7, phospho ATM, phospho ATR, phospho p53, Phospho Chk1, phospho Chk2, Bcl-2, phospho eif_2_α, eif_2_α, CHOP/GADD153, LC3B, phospho AMPKα, AMPK, phospho χH_2_AX, GRP78, PTEN, DDB2, pAKT, P21, P27, P62 ShRNA, ATG7 ShRNA, and secondary antibodies were purchased from Santa Cruz Biotechnologies (Santa Cruz, CA, USA). Fura 3 AM, DAPI, ER tracker, Acridine Orange, Ethidium Bromide, Salubrinal, Rapamycin, Chloroquine, Bafilomycin A1, Everolimus, GFP RFP LC3B Puncta kit from (Thermo Scientific). TUNNEL assay & CPD Elisa kits were procured from Abcam. Bradford reagent (catalog no. B6916; from Sigma-Aldrich. PVDF membrane was purchased from Bio-Rad, Hercules, CA. All other biochemicals used were of high purity biochemistry grade. All antibodies were purchased from Santa Cruz Biotechnology (USA) except Anti β-actin that was purchased from Sigma Aldrich.

### 2.2 Cell culture and UV-B exposure to HDFs

Human Dermal Fibroblasts (HDFs) were maintained in primary fibroblast expansion media from (HiMedia) supplemented with all the essentials including antibiotics, L-Glutamine, glucose (3.5 g/L), Hepes (15 mM), Penicillin (120mg/L), Streptomycin (270mg/L) and Fetal bovine serum (10% v/v) at 37 ºC in a humidified atmosphere of 5% CO_2_. Cells were exposed to UV-B using Daavlin UVA/UVB Research Irradiation Unit (Bryan, OH, USA) having digital control. The Lamps were maintained at a fixed distance of 24 cm from the surface of cell culture dishes. Majority of the resulting wavelengths (>90%) were in UV-B range (280–320 nm) because all the UV lamps are installed and adjusted in the irradiation unit in a series with UV-B lamps followed by UV-A and differentiated spatially from each other. UV-B irradiation of 10, 20, 30mJ/cm^2^ was used for initial standardization and dose optimization and 30mJ/cm^2^ dose was then selected and used for further experiments for mechanistic studies based on the analysis of cell toxicity induced by UV-B exposure to HDFs. Though 10mJ/cm^2^ is considered as the physiological dose mimicking the environmental dosage of UV-B in solar radiation spectrum [20], but it induces less cytotoxicity (10-20%) and shows least molecular changes to be selected for mechanistic studies. Before UV-B exposure, cell were first sensitized with chemical mediators like Rapamycin, Chloroquine, Salubrinal, Bafilomycin, P62 ShRNA, ATG7 ShRNA for a specified time period as per the particular experimental requirements to induce or inhibit autophagy. Cell mono-layers were then first washed with Dulbecco’s phosphate buffered saline (DPBS) and then UV-B -irradiated under a thin layer of pre-warmed DPBS. After irradiation, cells were again washed with DPBS twice and incubated in fresh medium with or without chemical mediators as per the experimental protocol requirements for 1, 3, 6 or 24h UV-B post -irradiation.

### 2.3 Cell viability Analysis

Colorimetric based MTT assay was employed for cell viability analysis as described earlier [21]. Briefly, the cells were seeded and incubated overnight in a humidified chamber. After treatment with chemicals/silencers or UV-B or both, the cells were further incubated for 24h. Cell viability was evaluated by assaying for the ability of functional mitochondria to catalyze the reduction of MTT to form formazan salt by an enzyme mitochondrial dehydrogenase that appear as crystals at the bottom of culture wells/dishes and was quantified by Multiskan Spectrum plate reader (Thermo Electron Corporation) at 570 nm using DMSO as detergent. Mean of 3 independent readings was taken for final quantification of data for result analysis.

### 2.4 Determination of Reactive oxygen species (ROS)

Dichlorofluorescin Diacetate (H_2_DCF-DA) staining was employed for the measurement of immediate Reactive Oxygen Species (ROS) generated upon UV-B exposure to HDFs (1h) post-irradiation, as described previously [20]. Briefly, HDFs were seeded in six-well plates and allowed to attach overnight. Cell monolayers were pre-treated with Salubrinal (25µM) and Everolimus (200nM) for 2h. Cells were after washed three times with DPBS and then exposed to UV-B 30 mJ/cm^2^. After UV-B irradiation, cells were then again washed with DPBS twice and incubated with fresh media with Salubrinal (25µM) and Everolimus (200nM) for 1h UV-B post-irradiation. After treatment, the cells were stained with 5 μM H_2_DCF-DA for 30 min at 37°C. The cells were then washed with DPBS thrice and observed immediately under a fluorescent microscope (Evos FL Color Imaging System from Thermo Scientific). Five random microscopic fields were selected and the intensity of fluorescence was quantified using the Image J software, as mentioned previously [22].

### 2.5 Confocal microscopy imaging of intracellular Ca^2+^

Ca^2+^ levels were determined by the Ca^2+^ indicator Fura 3 AM (Thermo Scientific) using confocal microscopy imaging as described previously. Briefly, after the cells were seeded to sterile cover slips and incubated overnight in humidified chamber to adhere. Cells were or weren’t treated with Salubrinal (25µM), Rapamycin (100nM), Chloroquine (50µM) and Bafilomycin A1 (100 nM) for 2h. Cells were then thrice washed with DPBS and exposed to UV-B treatment at 30 mJ/cm^2^ as described earlier and supplemented with fresh DMEM media with indicated concentrations of Salubrinal, Rapamycin, Chloroquine and Bafilomycin as required and incubated further for 6h post-UV-B –irradiation. HDFs were loaded with fluorescent Ca^2+^ indicator dye Fura 3 AM at 5 µM for 45 min before imaging post 6h UV-B –irradiation. Cells were washed three times with live cell imaging solution for imaging using a laser-scanning confocal microscope (Olympus Fluoview FV1000) by using a ×60 oil immersion objective lens. Five random microscopic fields were selected and the intensity of fluorescence was quantified using the Image J software, as mentioned previously.

### 2.6 siRNA-mediated knockdown of P62 and Atg7 expression

Validated P62 siRNA and Atg7 siRNA were purchased from Santa Cruz Biotechnology. siRNAs and lipofectamine (Invitrogen) were diluted into Opti-MEM I reduced serum medium (Invitrogen) per the manufacturer’s instructions. HDFs were incubated for 16h with transfection mixture at a final siRNA concentration of 50 pmol as described previously [23], after then exposed to UV-B (30 mJ/cm^2^) and were final supplemented with fresh medium for further 6h UV-B post –irradiation, as described earlier.

### 2.7 Protein isolation and Western blotting

Cells were trypsinized, harvested in PBS (pH 7.4), centrifuged, and resuspended in RIPA buffer (Sigma-Aldrich). After incubation for 45 min at 4°C, cell lysates were centrifuged at 17,530 *g* for 30 min at 4°C to remove cellular debris. Protein concentrations were determined by Bradford reagent. For Western blotting, 30–80 μg protein loads were denatured at 100°C for 3 min in Laemmli buffer. Protein samples were resolved on 4%– 15% SDS gels at 70-80 V. Proteins were electro transferred to PVDF membrane using Bio-Rad Mini Transblot Electrophoretic Transfer unit. Membranes were blocked in 5% fat free dry milk/ 3% BSA in 50 mM Tris, pH 8.0, with 150 mM sodium chloride, 2.6 mM KCl, and 0.05% Tween20 for 2h. Primary antibodies were used either in fat-free milk or BSA and incubated overnight at 4°C: anti-GRP78, anti-SQSTM1/p62, anti-Bcl_2_, anti-pmTOR, anti-mTOR, anti-CHOP/GAD153, anti-pelf_2_α, anti-elf_2_α, anti-BECN1, anti-ATG7, anti-pATM, anti-pATR, anti-pP53, anti-pChk1, anti-LC3B, anti-pAMPKα, anti-AMPKα, anti-pχH_2_AX, anti-PTEN, anti-pAKT (Cell Signaling Technology, Danvers, MA); anti-CPD from OxiSelect^TM^ UV-Induced DNA Damage staining kit (CPD quantification kit from Cell Bio Labs, Inc. San Diego, CA, USA) and mouse, anti-actin (Sigma-Aldrich). Goat anti-rabbit and goat anti-mouse immunoglobulin G antibodies conjugated with HRP (Santa Cruz Biotechnologies) were used as secondary antibodies. Chemiluminescence was detected by Immobilon chemiluminescent HRP substrate (EMD-Millipore, Billerica, MA) and visualized by Molecular Image ChemiDocTM XRS+ (Bio-Rad). Densitometric measurement of the bands was performed using Image LabTM software (version 3.0; Bio-Rad).

### 2.8 TUNEL assay

TUNEL assay was performed with an In Situ Direct DNA Fragmentation (TUNEL) Assay Kit (ab66108) from (Abcam) according to the manufacturer instructions as described previously [19]. Briefly, after the cells were seeded in dishes and incubated overnight in humidified chamber to adhere. Cells were or weren’t treated with Rapamycin (100nM) and Chloroquine (50µM) for 2h. Cell monolayer’s were after washed thrice with DPBS and exposed to UV-B (20 and 30 mJ/cm^2^). Again the cells were washed thrice with DPBS and supplemented with fresh DMEM media with or without indicated concentrations of Rapamycin (100nM) and Chloroquine (50µM) as required and incubated further for 6h post-UV-B –irradiation. Cell smears after fixation; blocking and permeabilization were incubated with TUNEL reaction mixture for 1h and wrapped in aluminium foil to avoid light exposure at 37°C and counterstained with RNase/PI solution for additional 20 minutes. Substrate solution was added and cells were imaged by a florescent microscope (Evos FL Colour Imaging System from Thermo Scientific) for detection of TUNNEL positive cells and the intensity of fluorescence was quantified using the Image J software.

### 2.9 ELISA based detection of Cyclobutane Pyrimidine dimers (CPD’s) and Pyrimide-(6-4)-Pyrimidone photoproducts (6, 4 PP’s)

UV-B induced CPD/6,4 PP adducts were quantified with an in situ OxiSelect^TM^ UV-Induced DNA Damage staining kit (CPD/6,4 PP quantification kit from Cell Bio Labs, Inc. San Diego, CA, USA) according to the manufacturer’s instruction. Briefly, after the cells were seeded and allowed to adhere overnight. Cells were or weren’t treated with Salubrinal (25µM), Rapamycin (100nM) and Chloroquine (50µM) for 2h or were silenced for P62 using P62 ShRNA as described previously. After the corresponding treatments to HDFs, cell monolayer’s were exposed to UV-B at (20 and 30 mJ/cm^2^) and supplemented with fresh DMEM media with or without indicated concentrations of Salubrinal (25µM), Rapamycin (100nM) and Chloroquine (50µM for further 6h post-UV-B –irradiation. DNA was isolated and incubated with the anti-CPD antibody for overnight on an orbital shaker at room temperature. Then the cells were washed and incubated with secondary FITC-conjugated antibody for 2h. The absorbance was measured at 450 nm using the Multiskan Spectrum plate reader (Thermo Electron Corporation). A mean of three independent readings was used to quantify the data for final result analysis.

### 2.10 Dual Acridine Orange/Ethidium Bromide (AO/EtBr) fluorescent staining for detection of DNA damage

Briefly, after the cells were seeded in 6 well plates and allowed to adhere overnight in incubator. Cells were left untreated or treated with Salubrinal (25µM), Rapamycin (100nM) and Chloroquine (50µM) for 2h and washed thrice with DPBS. Cells were then exposed to UV-B at 30mJ/cm^2^ and again washed thrice with DPBS. Cells were then supplemented with fresh DMEM media with or without indicated concentrations of Salubrinal, Rapamycin and Chloroquine as explained for additional 6h post-UV-B -irradiation. Dual fluorescent staining solution (1μl) containing 100 μg/ml AO and 100 μg/ml EtBr was added to cell monolayer’s for 5 minutes at RT and then covered with coverslip and were fixed as described previously [24]. The morphology of apoptotic cells were examined within 20 min using a fluorescent microscope (Evos FL Colour Imaging System from Thermo Scientific). Dual Acridine orange/Ethidium bromide (AO/EtBr) staining method was repeated at least 3 times for quantification and the intensity of fluorescence was quantified using the Image J software.

### 2.11 P62, LC3B, pP53, pχH_2_AX, PTEN, pAKT, DDB2, P21 and P27 immunostaining

Cultured cells were seeded on coverslips in six-well plates and incubated in the presence or absence of indicated concentrations of Salubrinal (25µM), Rapamycin (100nM) and Chloroquine (50µM) for 2h. Cells were then exposed to UV-B (30 mJ/cm^2^) and again washed thrice with DPBS and supplemented with fresh DMEM media with indicated concentrations of Salubrinal (25µM), Rapamycin (100nM) and Chloroquine (50µM) for required post UV-B time intervals of 1, 6 and 24h and were fixed in 4% paraformaldehyde for 15 min at room temperature. Cells were permeabilized in PBS with 0.1% TritonX-100 at room temperature for 10 min. Nonspecific binding sites were blocked by incubating the cells with 10% normal goat. Cells were incubated with P62, LC3B, pP53, pχH_2_AX, PTEN, pAKT, DDB2, P21 and P27 antibodies at a dilution of 1:100 in 0.1% Triton X-100 in PBS for overnight at 4 ºC, then washed and incubated with Alexa Fluor 488/594 conjugated antimouse or antirabbit secondary antibody as required at a dilution of 1:500 in DPBS for 2h at room temperature in dark conditions. Cells were then washed three times with PBS and stained with DAPI 1 μg/ml in PBS. The coverslips were mounted on glass slides, and cells were imaged by a laser-scanning confocal microscope (Olympus Fluoview FV1000) by using n ×60 oil immersion objective lens. Five random microscopic fields were selected and the intensity of fluorescence was quantified using the Image J software, as mentioned previously.

### 2.12 GFP-RFP-LC3B Puncta assay for detection of autophagy

The Premo^TM^ Autophagy Tandem Sensor RFP-GFP-LC3B puncta assay combines the ability to monitor the various stages of autophagy (through LC3B protein localization) with the high transduction efficiency and minimal toxicity of BacMam 2.0 expression technology. The protocol was adopted as per the manufacturer’s instructions and as described previously. For analysis of autophagy, simply after the cells were seeded in dishes and allowed to adhere overnight in incubator. BacMam 2.0 RFP-GFP-LC3B reagent was added to HDFs, incubated overnight to ensure maximum protein expression. The cells were thrice washed with DPBS. Then the cells were or weren’t treated with the Rapamycin (100nM) and Chloroquine (50µM) for 2h. The cells were then exposed to UV-B (30 mJ/cm^2^) and again washed thrice with DPBS and supplemented with fresh DMEM media with indicated concentrations of Rapamycin (100nM) and Chloroquine (50µM) for additional post UV-B time interval of 6h. Cells were then again washed with DPBS thrice and visualized using standard GFP (green fluorescent protein) and RFP (red fluorescent protein) settings. This tandem RFP-GFP sensor capitalizes on the pH difference between the acidic autolysosome and the neutral autophagosome and the pH sensitivity differences exhibited by GFP (green fluorescent protein) and RFP (red fluorescent protein) to monitor progression from the autophagosome to autolysosome. The puncta positive cells were imaged by a laser-scanning confocal microscope (Olympus Fluoview FV1000) by using n ×60 oil immersion objective lens. Five random microscopic fields were selected and the intensity of fluorescence was quantified using the Image J software, as mentioned previously.

### 2.13 Statistical analysis

Data is expressed as the mean ± standard deviation (SD). INSTAT statistical software was used to perform statistical analysis. Data represented is Mean• ±S.E. from three independent experiments. Comparisons between two groups were performed by student’s t-test and among groups by One-way ANOVA for statistical significance. p≤0.05, p<0.01, p<0.001 was considered as statistically significant.

## 3. Results

### 3.1 Improving autophagy levels and relieving ER stress response in HDFs reduces DNA photo adducts (CPD’s and 6,4 PP’s) in UV-B exposed HDFs

UV-B –irradiation to HDFs leads to formation of DNA photo-adducts (CPD’s and 6, 4 PP’s) in an intensity dependent manner in 6h UV-B post-irradiation, (^**^p<0.01 in UV-B 20mJ/cm^2^, ^#^p<0.001 in UV-B 30mJ/cm^2^ compared to control levels) (Fig. 1A). Improving autophagy levels in HDFs with Rapamycin (100nM) and upon relieving ER stress response with Salubrinal (25µM), an eif^2^α inhibitor in UV-B –irradiated HDFs significantly reduces the formation of both CPD’s and 6, 4 PP’s by 0.5 folds, (^**^p<0.01 in Salubrinal 25µM+UV-B 30mJ/cm^2^, ^**^p<0.01 in Rapamycin 100nM+UV-B 30mJ/cm^2^ compared to UV-B 30mJ/cm^2^ exposed only cells), (Fig. 1A). Inhibition of autophagy response with Chloroquine (50µM) has little effect on the formation of CPD’s whereas increases the formation of 6, 4 PP’s by about 0.2 folds in UV-B treated HDFs, (^#^p<0.001 in CHQ 50µM +UV-B 30mJ/cm^2^ compared to UV-B exposed only), (Fig 1A). ER stress response which is the immediate event after UV-B induced oxidative stress response upon –irradiation to cells normally leads to induction of autophagy to clear the damaged cellular debris. Here, we found that upon relieving ER stress response with Salubrinal (25µM) also reduces the DNA photo-adducts in UV-B treated HDFs in 6h UV-B post-irradiation. To confirm our details whether improving autophagy levels also improves the UV-B -induced photo-damage response in HDFs. We silenced key autophagy cargo protein P62 and found that it significantly impacts the formation of photo-adducts moreso CPD’s than that of 6, 4 PP’s and increases the formation of CPD’s by 0.5 folds and of 6, 4 PP’s by 0.4 folds, (^#^p<0.001 for CPD’s, ^*^p<0.5 for 6,4, PP’s in P62 ShRNA + UV-B 30mJ/cm^2^ treated compared to UV-B 30mJ/cm^2^ exposed only). P62 ShRNA only treated had negligible effect on the formation of DND photo-adducts compared to both control and UV-B treated cells, (^*^p<0.5 in P62 ShRNA treated only compared to control cells), (Fig. 1B). We confirmed the Elisa based results through Immunoflorescence as well through western blotting analysis and found that the expression levels of CPD’s are significantly increased in UV-B treated HDFs in an intensity dependent manner in 6h UV-B post-irradiation (Fig. 1D & 1E, Fig, 1F & 1G respectively). Inhibiting autophagy response with Chloroquine increases the expression of CPD’s in Immunoflorescence, (**p<0.01 compared to UV-B exposed only), (Fig. 1D & 1E), whereas Rapamycin treatment significantly reduces the formation of DNA photo-adducts in western analysis, (^**^p<0.01 compared to UV-B exposed only), (Fig. 1F & 1G). To check whether increasing autophagy levels in HDFs has any effect on the cellular viability in UV-B treated HDFs. We found that UV-B 30mJ/cm^2^ decreases the cell viability by 35% compared to control. Rapamycin treatment has no significant effect on restoring the cellular viability in UV-B treated HDFs at UV-B 30mJ/cm^2^ treatment whereas Chloroquine treatment significantly reduces the cell viability by 0.8 folds compared to UV-B only treated, (^*^p<0.05 for UV-B 30mJ/cm^2^ treated compared to control, ^*^p<0.05 for CHQ 50µM+ UV-B 30mJ/cm^2^ treated compared to UV-B exposed only), (Fig. 1C).

**Fig. 1:**
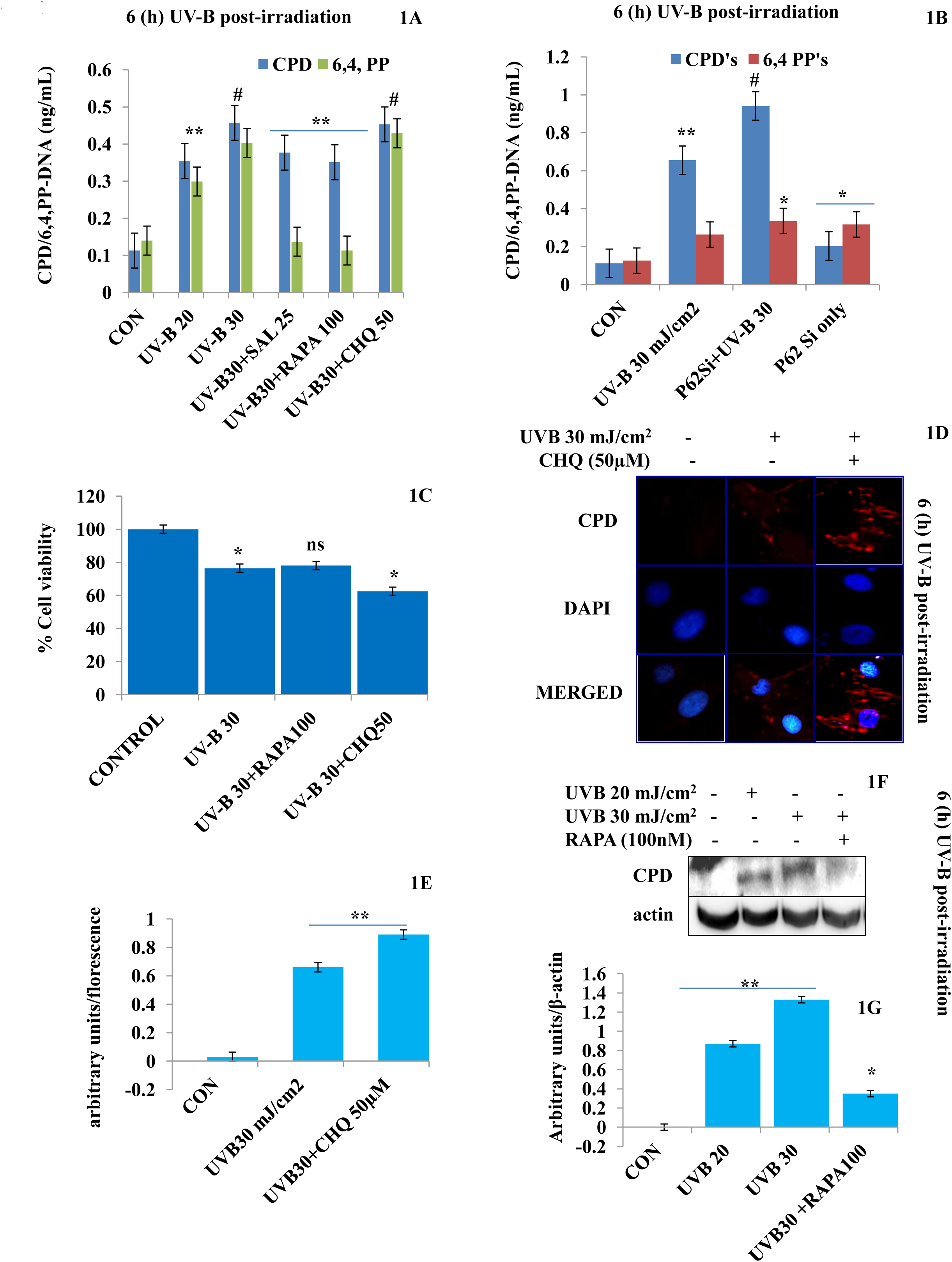
Improving autophagy levels and relieving ER stress response in UV-B exposed HDFs reduces DNA photo adducts (CPD’s and 6, 4 PP’s) (1A) ELISA based quantification of CPD’s and 6, 4, PP’s in UV-B 20-30mJ/cm^2^ exposed HDFs in 6h UV-B post-irradiation and effect of Salubrinal (25µM), Rapamycin (100nM) and Chloroquine (50µM) on UV-B –induced CPD and 6, 4 PP levels, (1B) ELISA based quantification of CPD’s and 6, 4, PP’s in P62 ShRNA treated HDFs, (1C) Cell viability analysis of UV-B 30mJ/cm^2^ exposed HDFs treated with Rapamycin (100nM) and Chloroquine (50 µM). (1D & 1E) Immunoflorescence analysis of CPD’s protein levels in microscopy in UV-B 30mJ/cm^2^ exposed HDFs treated with Chloroquine (50 µM). (1F & 1G) Western blotting analysis of CPD protein expression levels in HDFs in 6h UV-B post-irradiation. (*p≤ 0.05, **p≤ 0.01, #p≤ 0.001 were considered statistically significant).

### 3.2 Rapamycin treatment alleviates the UV-B induced TUNNEL positive as well as apoptotic cells in HDFs whereas Chloroquine treatment potentiates the effect of UV-B in florescent microscopy

Oxidatively induced DNA damage response is the hallmark of UV-B –induced photo damage in skin cells. To check whether improving autophagy levels in UV-B treated HDFs could alleviate the nuclear alterations. We performed TUNNEL assay and Acridine Orange/Ethidium Bromide co-staining. We found that UV-B treatment to HDFs induces TUNNEL positive cells in an intensity dependent manner in 6h UV-B post-irradiation. Rapamycin treatment (100nM) significantly reduces the florescence of TUNNEL positive cells in UV-B 30mJ/cm^2^ exposed HDFs by three folds whereas Chloroquine treatment (50µM) to HDFs increases the florescence of TUNNEL positive cells by 0.2 folds compared to UV-B 30mJ/cm^2^ only treated cells, (^**^p<0.01 for UV-B 20, UV-B 30 mJ/cm^2^ compared to control, (^*^p<0.05 for Rapamycin 100nM+ UV-B 30mJ/cm^2^, **p<0.01 for CHQ 50µM+UV-B 30mJ/cm^2^ treated compared to UV-B exposed only), (Fig. 2A & 2B). In AO/EtBr co-staining, we found that UV-B 30mJ/cm^2^ treatment increases the florescence of apoptotic nuclei in microscopic studies, citing nuclear damage, (^*^p<0.05 for UV-B 30 mJ/cm^2^ treated only compared to control). Salubrinal 25µM treatment significantly quenched the florescence of apoptotic nuclei by about 0.4 folds compared to UV-B only treated, (^*^p<0.05 for Salubrinal 25 µM+UV-B 30 mJ/cm^2^ treated compared to UV-B exposed only). Similarly, Rapamycin treatment also significantly quenches the florescence of apoptotic nuclei in microscopy compared to UV-B only treated by half, (^*^p<0.05 for Rapamycin 100 nM+UV-B 30 mJ/cm^2^ treated compared to UV-B exposed only). Chloroquine treatment to HDFs on the other hand, significantly increases the florescence of EtBr by 0.4 folds compared to UV-B only treated, (^**^p<0.01 for Chloroquine 50 µM+UV-B 30 mJ/cm^2^ treated compared to UV-B exposed only), (Fig. 2C & 2D).

**Fig. 2:**
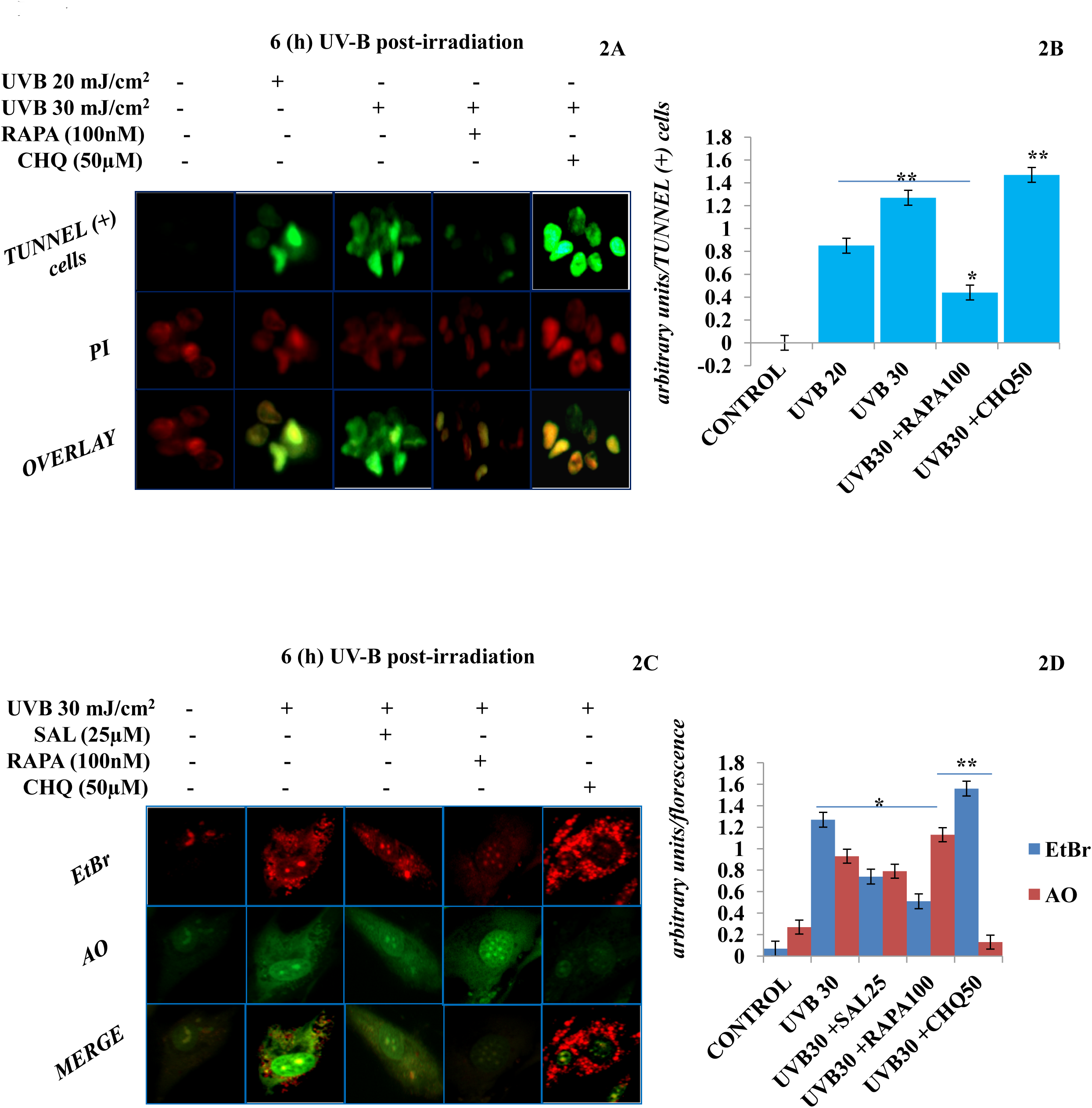
Autophagy promotion alleviates UV-B induced TUNNEL positive as well as apoptotic cells in HDFs in florescent microscopy. (2A & 2B) Florescent microscopic analysis of TUNNEL (+) cells in UV-B 30mJ/cm^2^ exposed HDFs showing nuclear damage in 6h UV-B post-irradiation and effect of Rapamycin (100nM) and Chloroquine (50µM) on UV-B –induced TUNNEL (+) cells. (2C & 2D) Acridine orange and Ethidium Bromide (AO-EtBr) co-staining depicting apoptotic (+) cells in UV-B 30mJ/cm^2^ exposed HDFs in 6h UV-B post-irradiation and effect of Salubrinal (25µM), Rapamycin (100nM) and Chloroquine (50µM) on UV-B –induced apoptosis. (*p≤ 0.05, **p≤ 0.01, were considered statistically significant).

### 3.3 UV-B exposure to HDFs induces immediate autophagy response (1 to 6h post-UV-B -irradiation) that vanishes in prolonged UV-B exposure (24h post UV-B –irradiation)

UV-B –irradiation to HDFs induces time dependent, i.e., immediate autophagy response (1 to 6h post-UV-B -irradiation) in an intensity dependent manner as is evident from the western blotting analysis of key autophagy marker proteins. UV-B exposure to HDFs modulates the expression profile of autophagy related proteins and increase the LC3BI to II lipidation, downregulates P62, upregulates pAMPKα but has in-significant effect on the expression levels of BECN1 indicating impaired autophagy upon UV-B exposure to HDFs, (^*^p<0.05 for LC3B compared to control, ^*^p<0.05 for P62 compared to control, ^*^p<0.05 for pAMPKα compared to control), (Fig. 3A, 3B, 3C, 3D & 3E respectively). We confirmed the induction of impaired autophagic flux response upon UV-B 30 mJ/cm^2^ exposure to HDFs through GFP-RFP-LC3B puncta assay which combines the ability to monitor the various stages of autophagy (through LC3B protein localization) and found that UV-B exposure to HDFs induces impaired flux that is evident from the lower GFP TO RFP conversion, although there are PUNCTA positive cells detected in UV-B exposed cells compared to control, (^*^p<0.05 for UV-B 30mJ/cm^2^ compared to control). Rapamycin treatment significantly rescued the UV-B exposed HDFs through improving autophagy levels as is clear from one fold improvement in conversion ratio of GFP to RFP depicting significant PUNCTA positive cells compared to UV-B exposed only, (^**^p<0.01 in UV-B 30mJ/cm^2^ compared to control cells). Chloroquine treatment to UV-B 30mJ/cm^2^ exposed HDFs showed least GFP-RFP-LC3B PUNCTA positive cells and has also stalled the conversion of GFP to RFP that depicts progression of autophagy and autophagosome maturation(Fig. 3F & 3G), (^*^p<0.05 for CHQ 50 µM+UV-B 30mJ/cm^2^ compared to UV-B exposed only). Further to confirm the western results, we checked the expression of autophagy marker protein in Immunoflorescence and found that the expression of P62 and LC3B lipidation from I to II is significantly modulated in 1h UV-B post –irradiation to HDFs, (^**^p<0.01 for P62 and LC3B in UV-B 30mJ/cm^2^ compared to control cells). Further we checked the expression of main DNA damage marker proteins during the same time interval (1h UV-B post-irradiation) and found that UV-B exposure significantly upregulates the expression of pχH2AX and Pp53 by 5 folds, (Fig. 3H & 3I), (^**^p<0.01 for UV-B 30mJ/cm^2^ compared to control). Primary Reactive Oxygen Species (ROS) are the main oxidative damage causing species in UV-B exposed cells. To check the effect of autophagy inducer Everolimus (200nM) and Salubrinal (25µM) which relieves ER in UV-B exposed cells on the generation of ROS species in UV-B exposed HDFs, we checked ROS species in UV-B exposed HDFs in 1h UV-B post –irradiation and found that primary ROS is developed immediately upon UV-B 30mJ/cm^2^ exposure to HDFs (1h), (^#^p<0.001 for UV-B 30mJ/cm^2^ compared to control). Salubrinal and Everolimus treatment to UV-B exposed HDFs significantly alleviates the production of ROS to half compared to UV-B levels, (^**^p<0.01 for SAL 25µM+UV-B 30mJ/cm^2^, **p<0.01 for Ever 200nM+UV-B 30mJ/cm^2^ compared to control), (Fig. 3J & 3K).

**Fig. 3:**
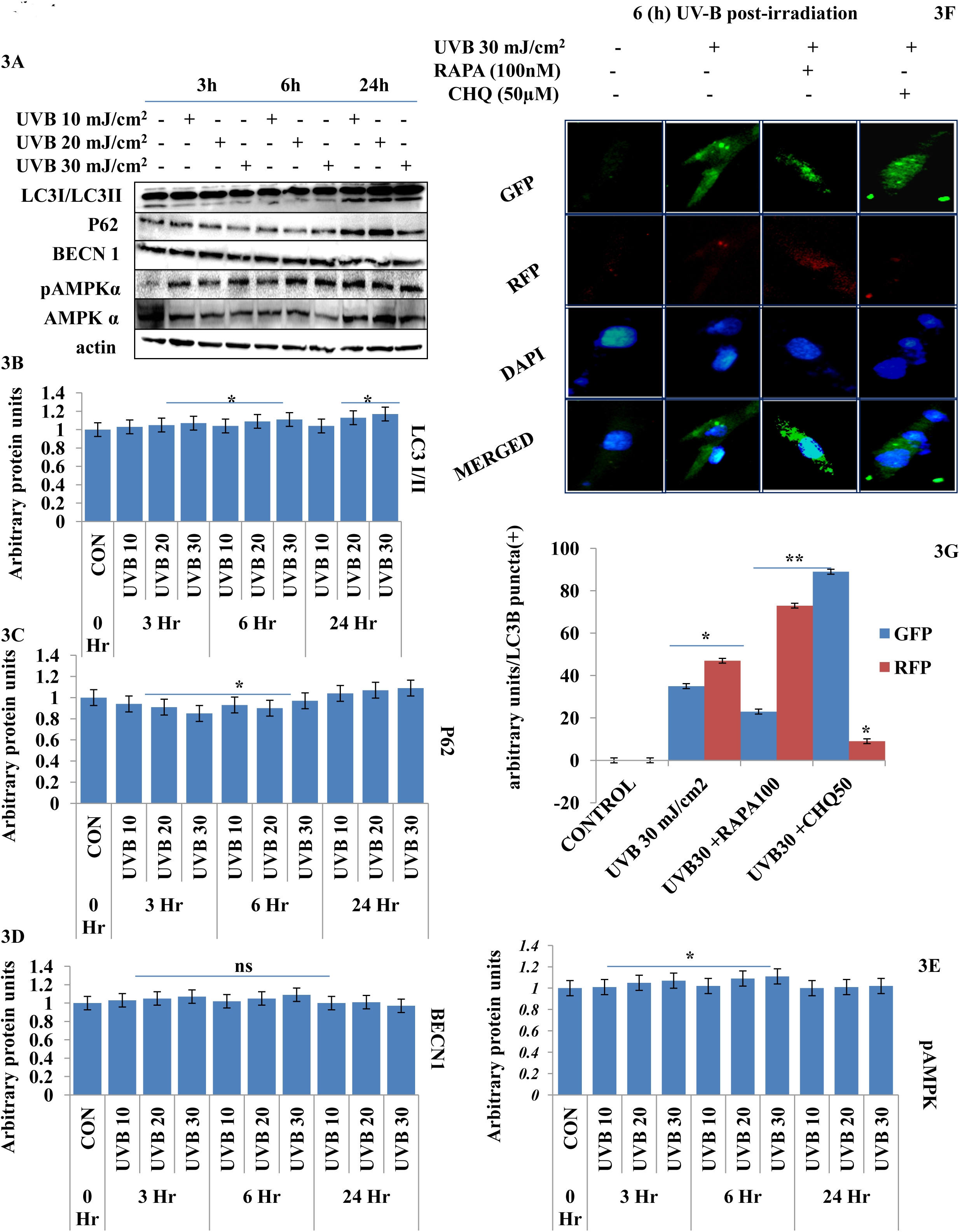
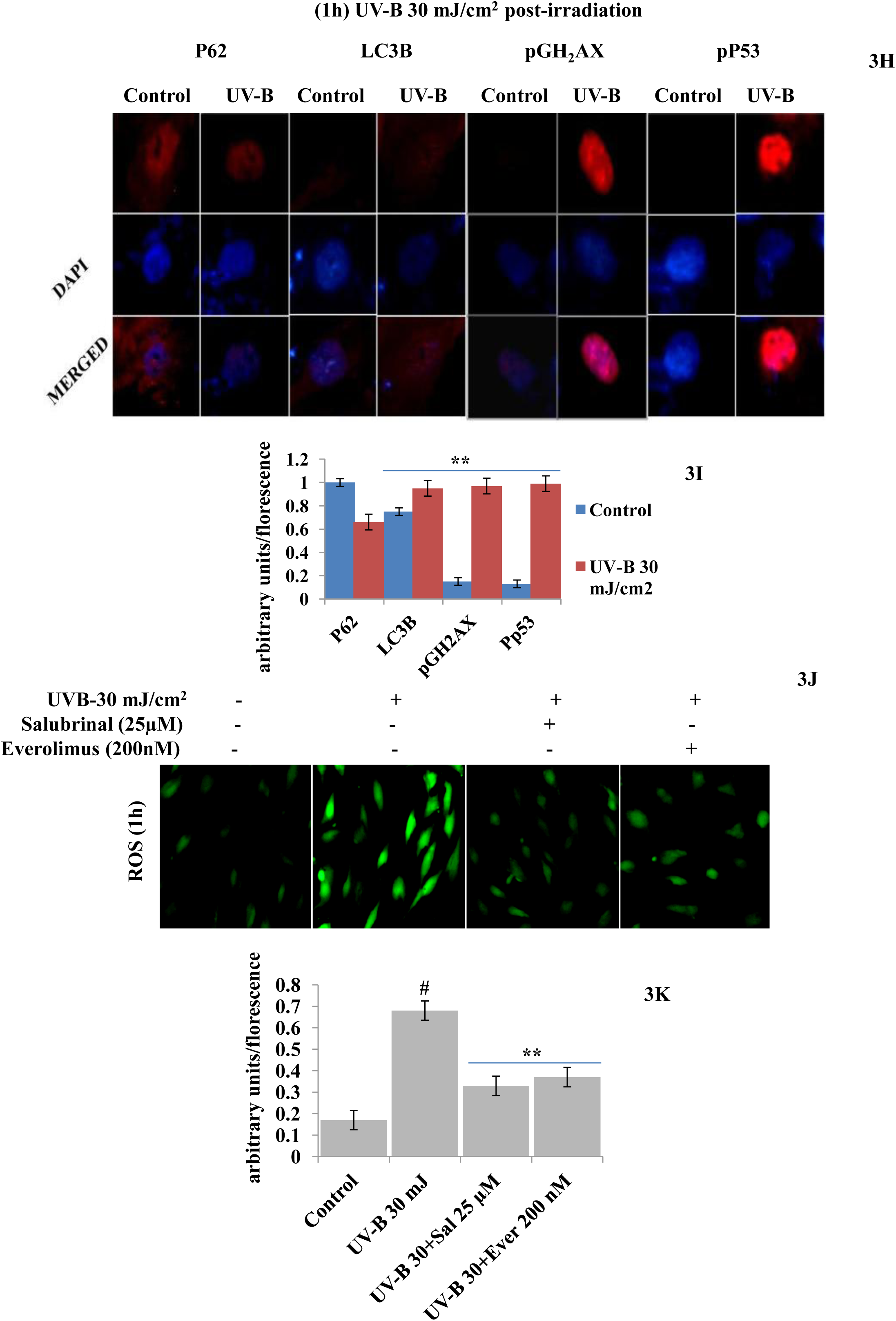
UV-B exposure to HDFs induces immediate autophagy response. (3A-3E) Western blotting analysis of key autophagy marker proteins showing time and intensity dependent response of autophagy in UV-B 10, 20 and 30mJ/cm^2^ exposed HDFs. (3F & 3G) GFP-RFP-LC3B PUNCTA assay for quantification of puncta (+) cells in UV-B 30mJ/cm^2^ exposed HDFs and effect of Rapamycin (100nM) and Chloroquine (50µM) on autophagy response in 6h UV-B post-irradiation. (3H & 3I) Immunoflorescence analysis of autophagy proteins P62 and LC3B and DNA damage marker proteins pχH2AX and pP53 in UV-B 30mJ/cm^2^ exposed HDFs in 1h UV-B post-irradiation in florescent microscopy. (3J & 3K) Reactive Oxygen Species (ROS) estimation in UV-B 30mJ/cm^2^ exposed HDFs treated with Salubrinal (25µM) and Everolimus (200nM) in 1h UV-B post-irradiation. (**p≤ 0.01, #p≤ 0.001, were considered statistically significant).

### 3.4 UV-B exposure to HDFs induces DNA damage response in time as well as in an intensity dependent manner. Improving autophagy response alleviates DNA Damage Response in UV-B –irradiated HDFs whereas blockage of autophagy potentiates the damage response

DNA damage response is the natural defense response to any genotoxic stimulus in cells and is attributed at repairing the damaged state to prevent tumorigenesis and to restore cellular homeostasis. Here we found that UV-B exposure at 10, 20 and 30mJ/cm^2^ induces DNA damage response in HDFs in a time and intensity dependent manner which was more aberrant in 30mJ/cm^2^ exposed HDFs as is evident from the western blotting analysis of DNA damage response proteins (Fig. 4A). The expression profile of key responsive proteins in DNA damage pathway were significantly modulated, pχH_2_AX (Fig. 4A & 4B), pATM (Fig. 4A & 4C), pATR (Fig. 4A & 4D) indicating damage response. Moreover, the expression level of pAKT, which has crucial role in pro-survival signaling and also inhibits apoptosis in UV-B response was also significantly increased and the increase was more profound in UV-B 30mJ/cm^2^ exposed HDFs, (Fig. 4A & 4E), (^*^p<0.5, ^**^p<0.01 for pχH2AX in UV-B exposed compared to control, ^*^p<0.5, for pATM in UV-B exposed compared to control, ^*^p<0.5, ^**^p<0.01 for pATR in UV-B exposed compared to control and ^*^p<0.5 for pAKT in UV-B exposed compared to control). As we were interested to study the mechanism of damage response and of that the damage as well the death was more prominent in prolonged UV-B exposure that is 24h UV-B post-irradiation. We confirmed our western results through Immunoflorescence in 6h UV-B post-irradiation to HDFs in confocal microscopy by imaging for the UV-B induced pχH2AX foci which directly reflect intensity and execution of damage response. In microscopic analysis, we found that UV-B 30mJ/cm^2^ exposure to HDFs induces the expression of pχH_2_AX foci, (^**^p<0.01 for pχH2AX in UV-B 30mJ/cm^2^ exposed compared to control). Salubrinal treatment (25µM) and Rapamycin (100nM) to UV-B exposed HDFs significantly rescued damage response in HDFs as is evident from the decreased expression of pχH2AX in confocal microscopy, (^*^p<0.05 for pχH2AX in SAL 25µM+UV-B 30mJ/cm^2^, RAPA 100nM+ UV-B 30mJ/cm^2^ treated compared to UV-B 30mJ/cm^2^ exposed only). Chloroquine (50µM) and Bafilomycin A1 (100nM) treatment significantly increased the expression level of pχH2AX nuclei in UV-B 30mJ/cm^2^ exposed HDFs citing autophagy positively regulates the damage responsive wing in UV-B exposed HDFs, (^**^p<0.01 for pχH2AX in CHQ 50µM+UV-B 30mJ/cm^2^, BAF A1 100nM+ UV-B 30mJ/cm^2^ treated compared to UV-B 30mJ/cm^2^ exposed only), (Fig. 4F & 4G). We further carried out the Immunoflorescence of DNA damage pathway protein DDB2 in confocal microscopy and found that DDB2 protein expression levels are significantly upregulated by 3 folds in UV-B 30mJ/cm^2^ exposed HDFs compared to control, (^*^p<0.05 in UV-B 30mJ/cm^2^ exposed compared to control). Rapamycin (100nM) significantly brought the DDB2 level to that of control whereas Chloroquine (50µM) treatment drastically increases the expression level of DDB2 protein further by 0.2 folds in UV-B exposed HDFs compared to UV-B only exposed, (^*^p<0.05 in RAPA 100nM+UV-B 30mJ/cm^2^ compared to UV-B only exposed, ^**^p<0.01 in CHQ 50µM+UV-B 30mJ/cm^2^ compared to UV-B exposed only), (Fig. 4H & 4I).

**Fig. 4:**
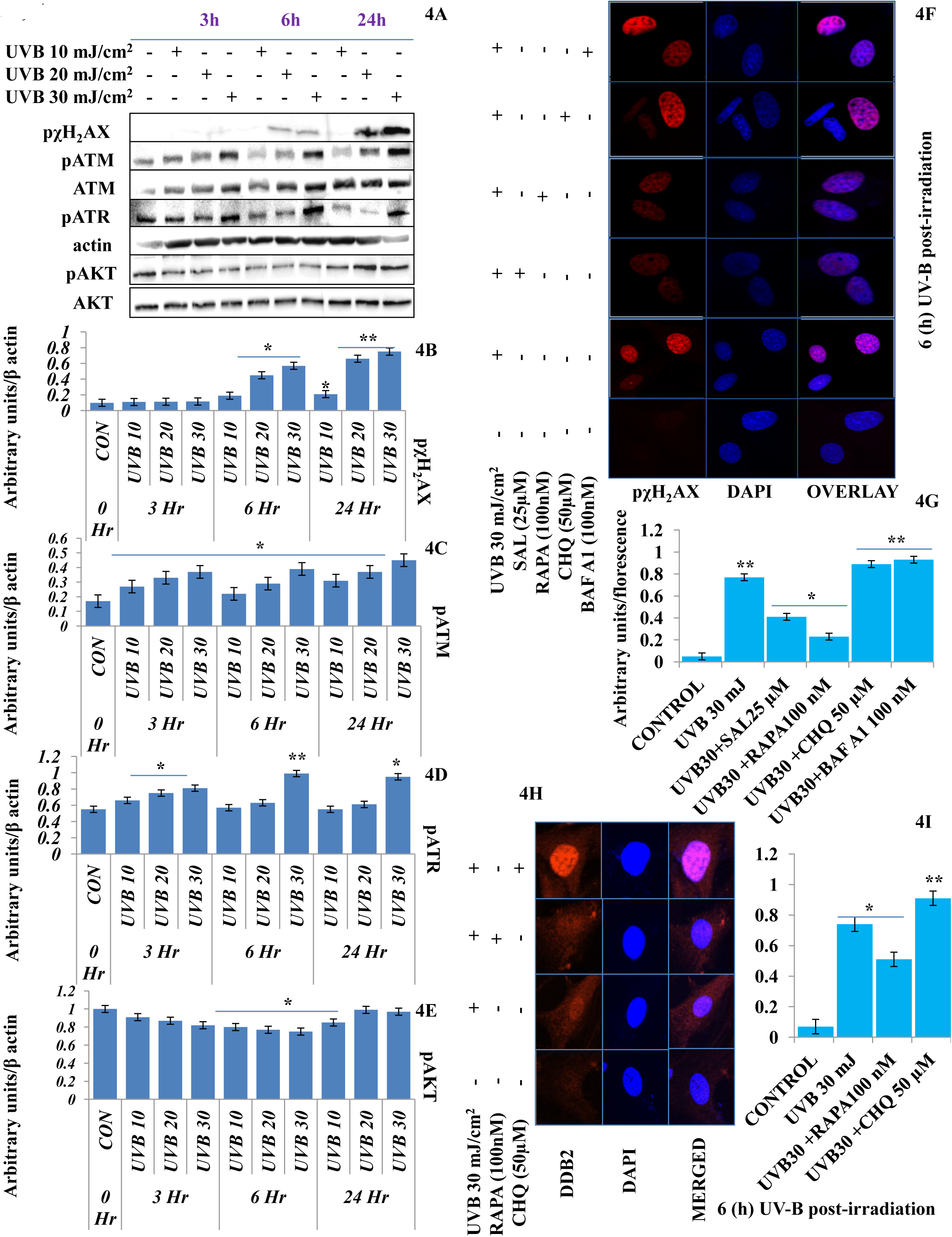
Improving autophagy response with Rapamycin alleviates DNA Damage in UV-B exposed HDFs. (4A-4E) Western blotting analysis of DNA damage response proteins showing time and intensity dependent effect of UV-B –irradiation on DNA damage response proteins in UV-B 10, 20, 30mJ/cm^2^ exposed HDFs. (4F & 4G) Immunoflorescence analysis of damage sensor protein pχH2AX foci in UV-B 30mJ/cm^2^ exposed HDFs in 6h UV-B post-irradiation and effect of Salubrinal (25µM), Rapamycin (100nM), Chloroquine (50µM) and Bafilomycin A1 (100nM) on pχH2AX expression levels under UV-B exposure to HDFs. (4H & 4I) Immunoflorescence analysis of DDB2 protein expression levels in UV-B mJ/cm^2^ exposed HDFs and effect of Rapamycin (100nM) and Chloroquine (50µM) on the expression levels of DDB2. (*p≤ 0.05, **p≤ 0.01, were considered statistically significant).

### 3.5 Salubrinal, (an eif_2_α inhibitor) relieves autophagy, alleviates DNA damage and prevents immediate ER calcium leakage from UV-B exposed HDFs

ER stress response is the immediate manifestation of oxidative stress response in UV-B exposure to HDFs. Here we used Salubrinal which is an eif_2_α inhibitor and relieves ER stress response upon UV-B exposure to cells. We first performed the cell viability assay of Salubrinal treated HDFs in UV-B exposed HDFs to find out if it imparts any cytotoxicity on its use in HDFs. UV-B exposure decreases the cellular viability in HDFs in an intensity dependent manner in 24h MTT assay, (Fig 5A). UV-B decreases the cellular viability of HDFs in MTT assay by 20%, 27% and 35% at 10, 20 and 30mJ/cm^2^ exposure, (^**^p<0.01 in UV-B 10, 20 and 30mJ/cm^2^ exposed compared to control). Salubrinal treatment at 10, 20 and 30 µM improved the cell viability by 1 folds, 0.5 folds and 0.5 folds in UVB 10mJ/cm^2^+ SAL 10µM, 20mJ/cm^2^+ SAL 20µM and UVB 30mJ/cm^2^+SAL 30µM treated HDFs respectively, (^*^p<0.05 UVB 10mJ/cm^2^+ SAL 10µM, 20mJ/cm^2^+ SAL 20µM and UVB 30mJ/cm^2^+SAL 30µM treated HDFs compared to UV-B 10, 20 and 30mJ/cm^2^ exposed only, respectively). It was found the Salubrinal treatment is safe upon 30µM in its use in HDFs. Further we checked the expression of autophagy, ER stress response and DNA damage response pχH_2_AX protein levels in UV-B exposed HDFs in western blotting in 24h UV-B post-irradiation upon Salubrinal treatment and found that although UV-B exposure induces impaired autophagy flux response as is evident from the expression levels of P62, (Fig. 5B & 5C) and BECN1 (Fig. 5B & 5D). UV-B exposure to HDFs also induces ER stress response as evident from the expression of peif_2_α, (Fig. 5B & 5E) and also increases the expression of DNA damage sensor protein pχH_2_AX in western blotting in 24h UV-B exposure (Fig. 5B & 5F) but Salubrinal at (20µM) failed to significantly improve the ER stress and damage response events in UV-B exposed HDFs in 24h post-irradiation. We reduced the UV-B post -exposure time interval to 6h and further increased the working concentration of Salubrinal to 25µM, because both autophagy and ER stress response are the initial time interval as already reported in our previous study [19]. We found that ER stress induced upon UV-B exposure to HDFs in an intensity dependent manner in 6h UV-B post-irradiation was significantly relieved by treatment with Salubrinal 25µM as evident from the expression of key ER stress response proteins, peif_2_α, GRP78 and CHOP/GAD153, (^*^p<0.05, ^**^p<0.01 for peif_2_α, GRP78 and CHOP/GAD153 in SAL 25µM+UVB 30mJ/cm^2^ treated HDFs compared to UV-B 30mJ/cm^2^ exposed only). Rapamycin (100nM) treatment has also significantly relieved the UV-B induced ER stress response whereas Chloroquine (50µM) has failed to rescue the cells from stress response in UV-B exposed HDFs, (Fig. 5G & 5K). Salubrinal treatment to UV-B exposed HDFs has also reduced the autophagy response due to relieving of ER stress response as evident from the western blotting analysis of key autophagy markers P62 and BECN1, (^*^p<0.05, ^**^p<0.01 for P62 and BECN1 in SAL 25µM+UVB 30mJ/cm^2^ treated HDFs compared to UV-B 30mJ/cm^2^ exposed only), (Fig. 5G & 5K). Further, to check whether upon relieving ER stress response with Salubrinal has any impact on rescuing the UV-B 30mJ/cm^2^ exposed HDFs from aberrant DNA damage response? We checked for the expression of DNA damage sensor protein pχH_2_AX and pChk1 and found that Salubrinal (25µM) significantly reduces the expression of both pχH2AX and pChk1 in western blotting, (^*^p<0.05, ^**^p<0.01 for pχH2AX and pChk1 in SAL 25µM+UVB 30mJ/cm^2^ treated HDFs compared to UV-B 30mJ/cm^2^ exposed only) (Fig. 5H & 5L). Salubrinal (25µM) treatment has been able to significantly restore the expression levels of ant-apoptotic protein Bcl_2_, whose expression is dwindled upon UV-B 30mJ/cm^2^ exposure to HDFs, (^**^p<0.01 for Bcl_2_ in SAL 20µM+UVB 30mJ/cm^2^ treated HDFs compared to UV-B 30mJ/cm^2^ exposed only). Rapamycin (100nM) treatment has also restored the Bcl_2_ to a significant level whereas Chloroquine (50µM) treatment has failed to do so, (^*^p<0.05 for Bcl_2_ in RAPA 100nM+UVB 30mJ/cm^2^, ^**^p<0.01 in CHQ 50µM+UVB 30mJ/cm^2^treated HDFs compared to UV-B 30mJ/cm^2^ exposed only), (Fig. 5H & 5L). ER calcium leakage/depletion is the immediate event following UV-B exposure to HDFs, (^**^p<0.01 for UVB 30mJ/cm^2^ exposed HDFs compared to control). Salubrinal (25µM) treatment significantly prevents the ER Calcium leakage in confocal microscopy, (^#^p<0.001 for Salubrinal 25µM+UVB 30mJ/cm^2^ exposed HDFs compared to UV-B 30mJ/cm^2^ exposed only). Rapamycin (100nM) also alleviates UV-B exposed HDFs from Calcium depletion state whereas Chloroquine (50µM) and Bafilomycin A1 (100nM) potentiates the calcium depletion from ER as evident in confocal microcopy analysis with Fluo-3 staining, (^#^p<0.001 for Rapamycin 100nM+UVB 30mJ/cm^2^ exposed HDFs compared to UV-B 30mJ/cm^2^ exposed only, ^**^p<0.01 for Chloroquine 50µM+UVB 30mJ/cm^2^ exposed HDFs compared to UV-B 30mJ/cm^2^ exposed only, ^#^p<0.001 for Bafilomycin A1 100nM+UVB 30mJ/cm^2^ exposed HDFs compared to UV-B 30mJ/cm^2^ exposed only), (Fig. 5I & 5J).

**Fig. 5:**
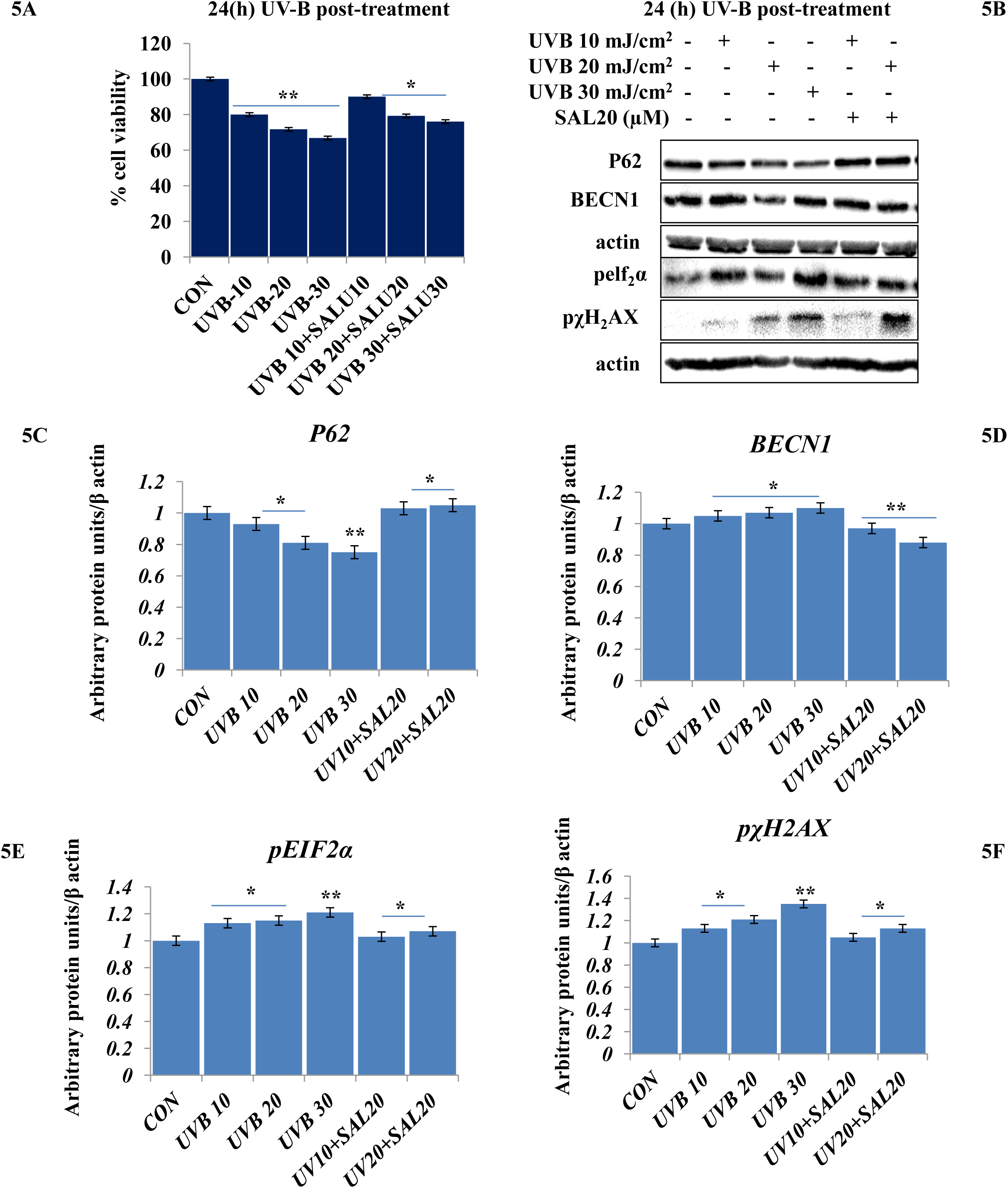
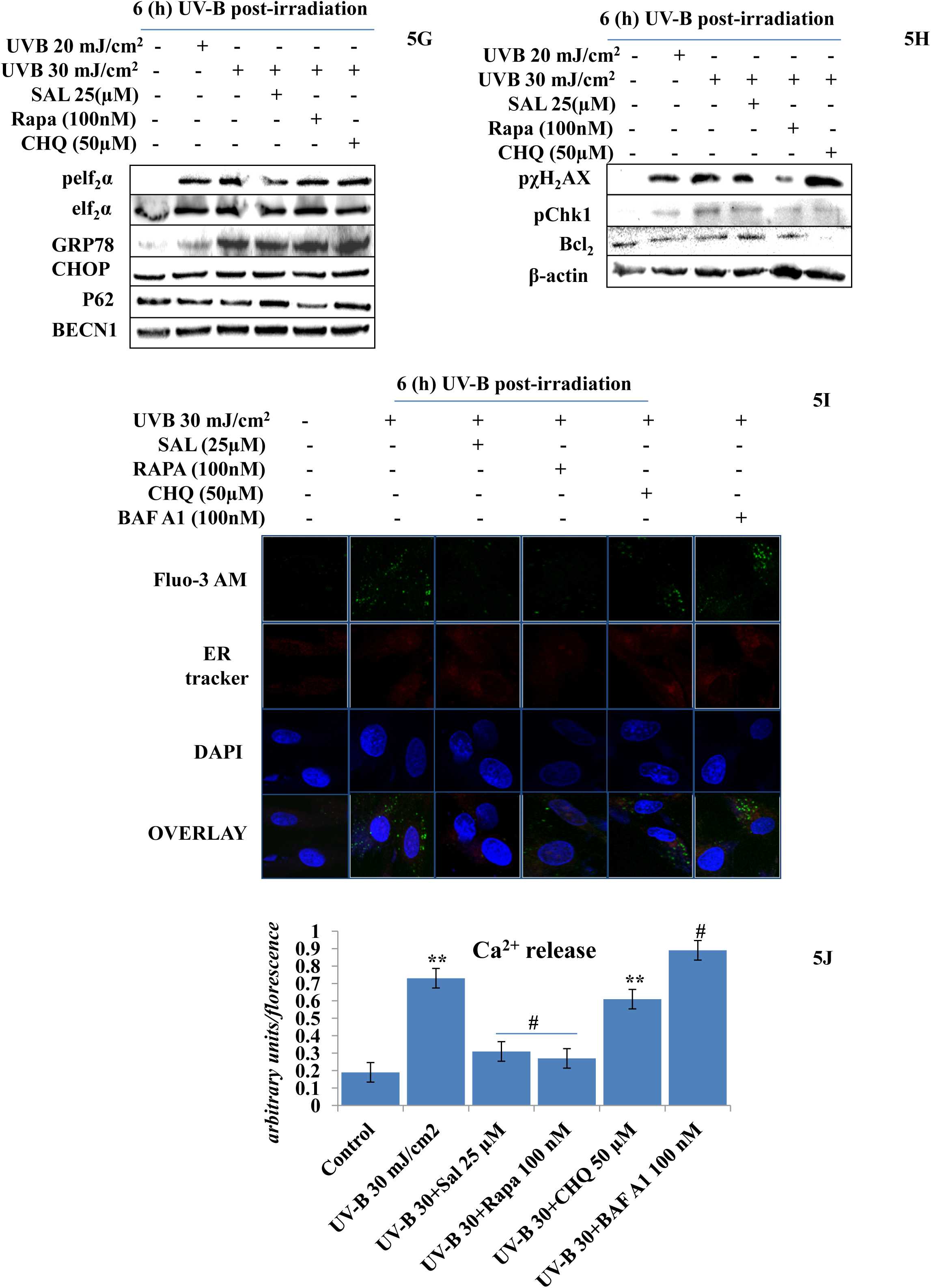
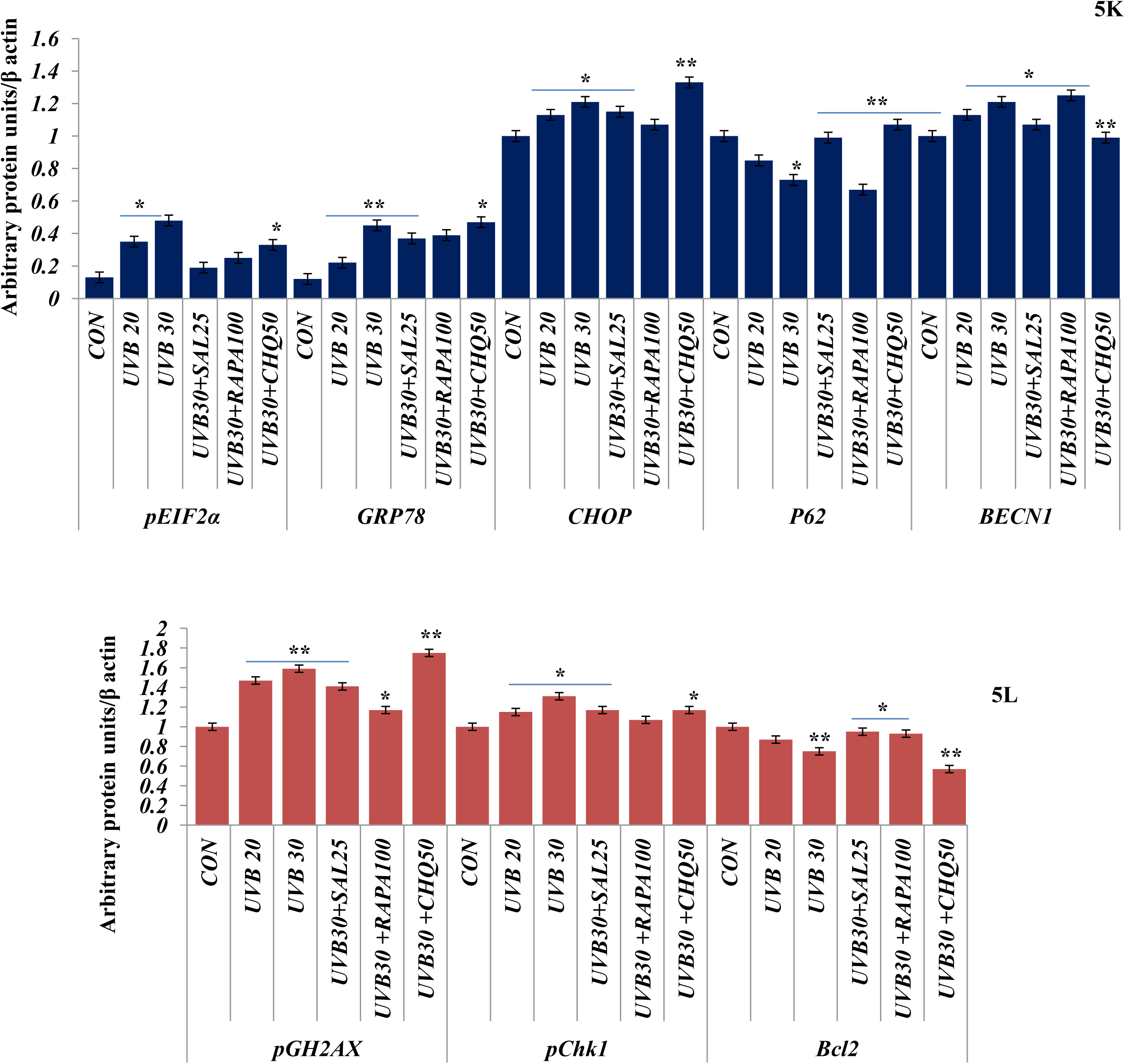
Salubrinal, (an eif_2_α inhibitor) alleviates DNA damage and prevents immediate ER calcium leakage from UV-B exposed HDFs. (5A) cell viability analysis of UV-B 10, 20, 30mJ/cm^2^ exposed HDFs in presence of Salubrinal (10, 20, 30µM) in 24h UV-B post-irradiation. (5B-5F) Western blotting analysis of ER stress, autophagy and DNA damage response proteins in 10, 20, 30mJ/cm^2^ exposed HDFs in 24h UV-B post-irradiation in presence of Salubrinal (20µM). (5G & 5K, 5H & 5L) Western blotting analysis of ER stress, autophagy and DNA damage response proteins respectively, in UV-B 20 and 30mJ/cm^2^ exposed HDFs in 6h UV-B post-irradiation in presence of Salubrinal (25µM), Rapamycin (100nM) and Chloroquine (50µM). (5I & 5J) Confocal microscopy analysis of ER calcium depletion in 30mJ/cm^2^ exposed HDFs in 6h UV-B post-irradiation in presence of Salubrinal (25µM), Rapamycin (100nM), Chloroquine (50µM) and Bafilomycin A1 (100nM). (*p≤ 0.05, **p≤ 0.01, ^#^p≤ 0.001 were considered statistically significant).

### 3.6 Autophagy blockage via P62 silencing potentiates DNA damage response in UV-B exposed HDFs

Autophagy is the main pathway that is activated in stress response in order to restore the normal homeostatic state of cells subjected to genotoxic stress. We blocked autophagy response in UV-B exposed HDFs through silencing autophagy cargo protein P62 to find the impact of autophagy blockage on DNA damage response upon UVB 30mJ/cm^2^ exposure to HDFs in 6h UV-B post-irradiation. Western blotting analysis confirmed the silencing of P62 and depicted 90% silencing efficiency, (^*^p<0.05 for P62 in P62 ShRNA treated HDFs compared to control). UVB 30mJ/cm^2^ exposure to HDFs showed mild autophagy induction as evident from the downregulation of P62 protein levels by 0.25 folds in western blotting analysis, (Fig. 6A & 6B), (^*^p<0.05 for P62 in UVB 30mJ/cm^2^ exposed HDFs compared to control). We found that P62 silenced HDFs show enhanced DNA damage response upon UVB 30mJ/cm^2^ exposure in 6h UV-B post-irradiation as is clear in western blotting analysis of DNA damage response proteins pχH_2_AX, pATR, pChk2, pP53, (Fig. 6A & 6B). P62 silencing enhanced the expression levels of pχH2AX, pATR, pChk2, pP53 by 1 fold in western blotting, (^#^p<0.001 for pχH_2_AX in P62 ShRNA+UVB 30mJ/cm^2^ treated HDFs compared to UV-B 30mJ/cm^2^ exposed only, ^**^p<0.01 for pATR in P62 ShRNA+UVB 30mJ/cm^2^ treated HDFs compared to UV-B 30mJ/cm^2^ exposed only, ^**^p<0.01 for pChk2 in P62 ShRNA+UVB 30mJ/cm^2^ treated HDFs compared to UV-B 30mJ/cm^2^ exposed only, ^*^p<0.05 for pP53 in P62 ShRNA+UVB 30mJ/cm^2^ treated HDFs compared to UV-B 30mJ/cm^2^ exposed only). P62 ShRNA only treated cells show negligible effect on the change in expression of DNA damage response proteins in western blotting analysis compared to control levels. We confirmed our western blotting results through Immunoflorescence in confocal microscopy by looking for the pχH_2_AX foci and found that UV-B 30mJ/cm^2^ induced pχH_2_AX foci are significantly increased in P62 silenced cells, (Fig. 6C & 6D), (^*^p<0.05 in UVB 30mJ/cm^2^ exposed HDFs compared to control, ^**^p<0.01 in P62 ShRNA+UVB 30mJ/cm^2^ treated compared to UV-B 30mJ/cm^2^ exposed only). Rapamycin (100nM) treatment to P62 silenced HDFs upon UV-B exposure decreased the pχH_2_AX foci by 2 folds in Immunoflorescence, (^*^p<0.05 in P62 ShRNA+UVB 30mJ/cm^2^+RAPA (100nM) treated compared to UVB 30mJ/cm^2^ exposed only). Chloroquine (50µM) treatment to P62 silenced cells upon UV-B exposure failed to alleviate the DNA damage response in HDFs as evident in Immunoflorescence of pχH_2_AX, (^**^p<0.01 in P62 ShRNA+UVB 30mJ/cm^2^+CHQ (50µM) treated compared to UVB 30mJ/cm^2^ exposed only), (Fig. 6C & 6D).

**Fig. 6:**
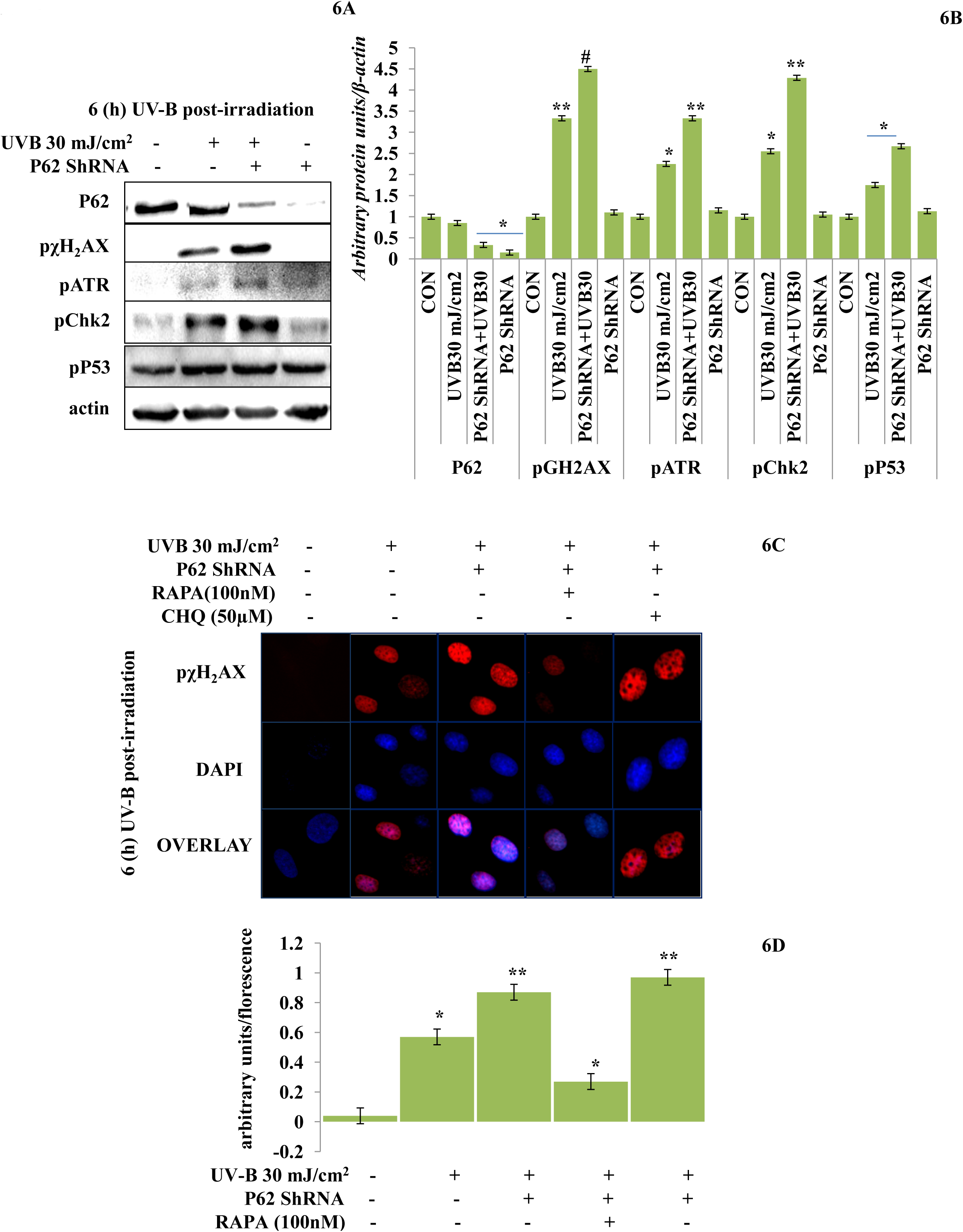
P62 silencing enhances DNA damage response in UV-B exposed HDFs. (6A & 6B) Western blotting analysis of DNA damage response proteins in P62 ShRNA treated HDFs in 6h UV-B 30mJ/cm^2^ post-irradiation. (6C & 6D) Immunoflorescence analysis of expression profile of DNA damage sensor protein pχH2AX in P62 ShRNA treated HDFs in 6h UV-B post-irradiation in presence of Rapamycin (100nM) and Chloroquine (50µM). (*p≤ 0.05, **p≤ 0.01, ^#^p≤ 0.001 were considered statistically significant).

### 3.7 Autophagy blockage via P62 silencing dwindles the tumour suppressor PTEN/AKT pathway in UV-B exposed HDFs

PTEN/AKT pathway is the main tumour suppressor pathway that promotes cell survival and reduces tumorigenesis in UV-B induced photo-damage to HDFs through regulation of autophagy response. Here we checked whether blockage of autophagy has any substantial impact on the PTEN/AKT pathway in UV-B 30mJ/cm^2^ exposed HDFs in 6h UV-B post-irradiation. We found that UV-B –irradiation to HDFs significantly downregulates the expression level of PTEN by 0.2 folds and increases the expression of pAKT protein by 0.2 folds in western blotting analysis, (Fig. 7A & 7B), (^*^p<0.05 for PTEN in UVB 30mJ/cm^2^ exposed compared to control, ^**^p<0.01 for pAKT in UVB 30mJ/cm^2^ exposed compared to control levels). Autophagy blockage with P62 silencing significantly downregulated the expression of PTEN levels in UV-B exposed HDFs further by 1 fold compared to UV-B exposed HDFs, (*p<0.05 for PTEN in P62 ShRNA + UV-B 30mJ/cm^2^ treated compared to UVB 30mJ/cm^2^ exposed only). We got very contrasting results for pAKT as the protein expression level of pAKT were upregulated by 0.2 folds in P62 silenced cells compared to UV-B exposed cells, (**p<0.01 for pAKT in P62 ShRNA + UV-B 30mJ/cm^2^ treated compared to UVB 30mJ/cm^2^ exposed only). We got similar results for PTEN and pAKT in Immunoflorescence in confocal microscopy. PTEN protein expression was downregulated upon UV-B exposure to HDFs in Immunoflorescence, (*p<0.05 for PTEN in UVB 30mJ/cm^2^ exposed compared to control). Autophagy activator Rapamycin (100nM) significantly restores the PTEN expression level by 2 folds whereas, inhibitor of autophagy Chloroquine (50µM) treatment to UV-B exposed HDFs could not restore the PTEN expression but adversely decreases the expression level of PTEN by 1 fold compared to UV-B only exposed as evident in Immunoflorescence, (Fig. 7C & 7D), (*p<0.05 in UV-B 30mJ/cm^2^+RAPA 100nM treated compared to UVB 30mJ/cm^2^ exposed only, **p<0.01 in UV-B 30mJ/cm^2^+ CHQ 50µM treated compared to UVB 30mJ/cm^2^ exposed only). pAKT expression level in UV-B exposed HDFs on the other hand is upregulated by 2 folds in Immunoflorescence compared to control levels (Fig. 7E & 7F), (*p<0.05 in UV-B 30mJ/cm^2^ exposed compared to control). Rapamycin (100nM) treatment significantly brought the expression of pAKT to control levels whereas Chloroquine (50µM) drastically increased the expression in UV-B exposed HDFs by 0.2 folds compared to UV-B exposed only, (*p<0.05 in UV-B 30mJ/cm^2^+ RAPA 100nM treated compared to UVB 30mJ/cm^2^ exposed only, **p<0.01 in UV-B 30mJ/cm^2^+ CHQ 50µM treated compared to UVB 30mJ/cm^2^ exposed only).

**Fig. 7:**
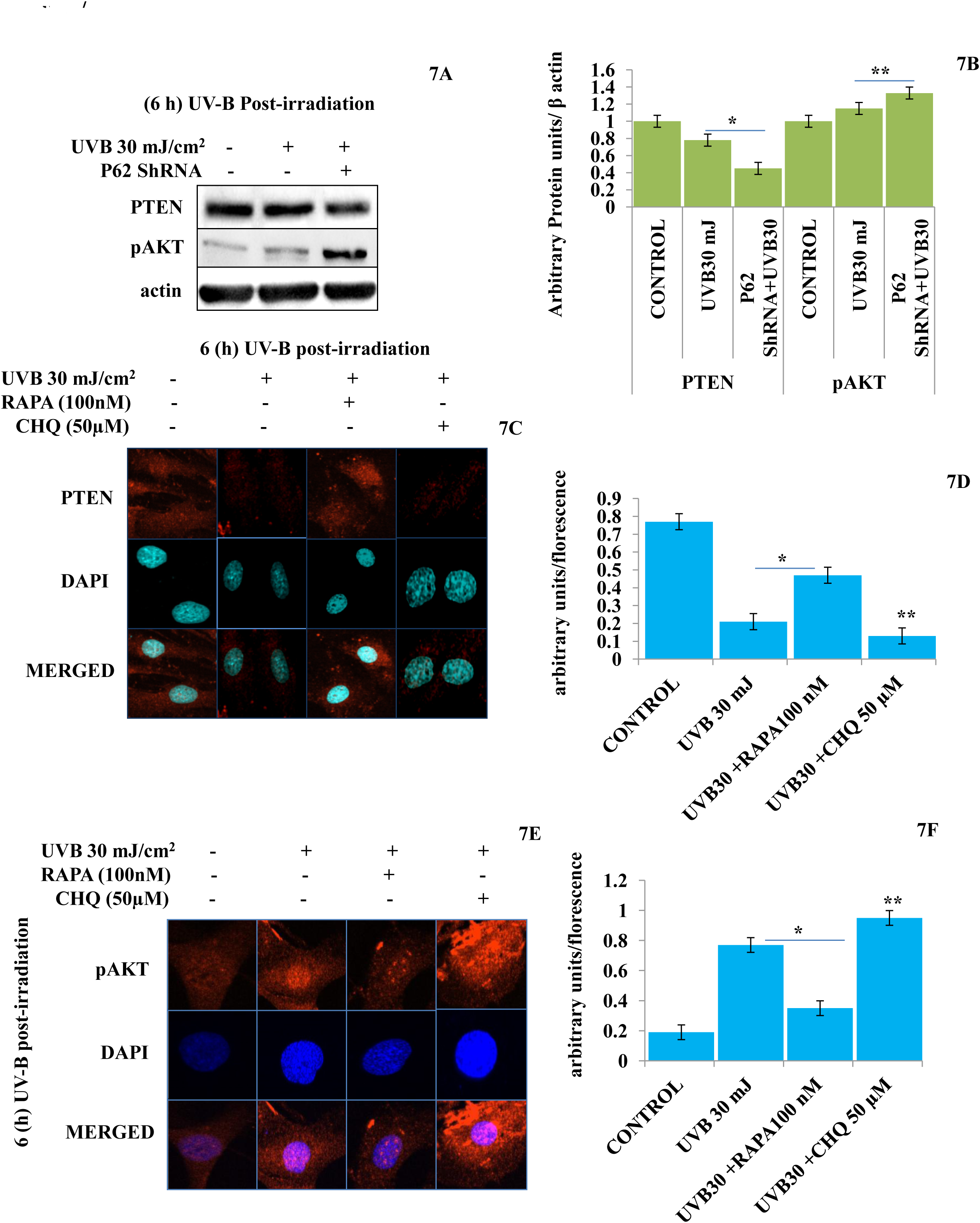
P62 silencing dwindles the tumour suppressor PTEN/AKT pathway in UV-B exposed HDFs. (7A & 7B) Western blotting analysis of PTEN/pAKT protein expression levels in P62 ShRNA treated HDFs in 6h UV-B 30 mJ/cm^2^ post-irradiation. (7C & 7D) Immunoflorescence analysis of PTEN protein expression levels in UV-B 30 mJ/cm^2^ exposed HDFs in 6h UV-B post-irradiation in presence of Rapamycin (100nM) and Chloroquine (50µM). (7E & 7F) Immunoflorescence analysis of pAKT protein expression levels in UV-B 30 mJ/cm^2^ exposed HDFs in 6h UV-B post-irradiation in presence of Rapamycin (100nM) and Chloroquine (50µM). (*p≤ 0.05, **p≤ 0.01, were considered statistically significant).

### 3.8 Rapamycin improves cell cycle regulation by regulating the expression of cell cycle regulatory proteins in Immunoflorescence in UV-B exposed HDFs

Cell cycle regulatory proteins play an important role in quality control and respond to any genotoxic insult and thus prevent cancer development in cells. In line with this, we checked the effect of chemically stimulated autophagy response on main cell cycle regulator proteins in Immunoflorescence through confocal microscopy in 24h UV-B post-irradiation to HDFs. We found that the expression of P21 protein is upregulated in UV-B exposed HDFs by 3 folds compared to control levels, (**p<0.01 in UV-B 30mJ/cm^2^ exposed compared to control). Rapamycin (100nM) treatment improved the expression level of P21 to that of control, (*p<0.05 in UV-B 30mJ/cm^2^+ RAPA 100nM treated compared to UVB 30mJ/cm^2^ exposed only), whereas Chloroquine (50µM) drastically but significantly increased the expression of P21 in Immunoflorescence by 0.2 folds compared to UV-B only exposed levels, (**p<0.01 in UV-B 30mJ/cm^2^+ CHQ 50µM treated compared to UVB 30mJ/cm^2^ exposed only) (Fig. 8A & 8B). Similarly, we got higher expression in the protein levels of P27 in UV-B 30mJ/cm^2^ exposed HDFs by 3 folds compared to control levels, (**p<0.01 in UV-B 30mJ/cm^2^ compared to control). Salubrinal (25µM) and Rapamycin (100nM) treatment brought the expression of P27 to that of control levels, (*p<0.05 in UV-B 30mJ/cm^2^+ SAL 25µM treated compared to UVB 30mJ/cm^2^ exposed only, *p<0.05 in UV-B 30mJ/cm^2^+ RAPA 100nM treated compared to UVB 30mJ/cm^2^ exposed only), whereas Chloroquine (50µM) significantly increased the expression of p27 by 0.2 folds compared to UV-B levels in Immunoflorescence in UV-B exposed HDFs, (**p<0.01 in UV-B 30mJ/cm^2^ + CHQ 50µM treated compared to UVB 30mJ/cm^2^ exposed only), (Fig. 8C & 8D).

**Fig. 8:**
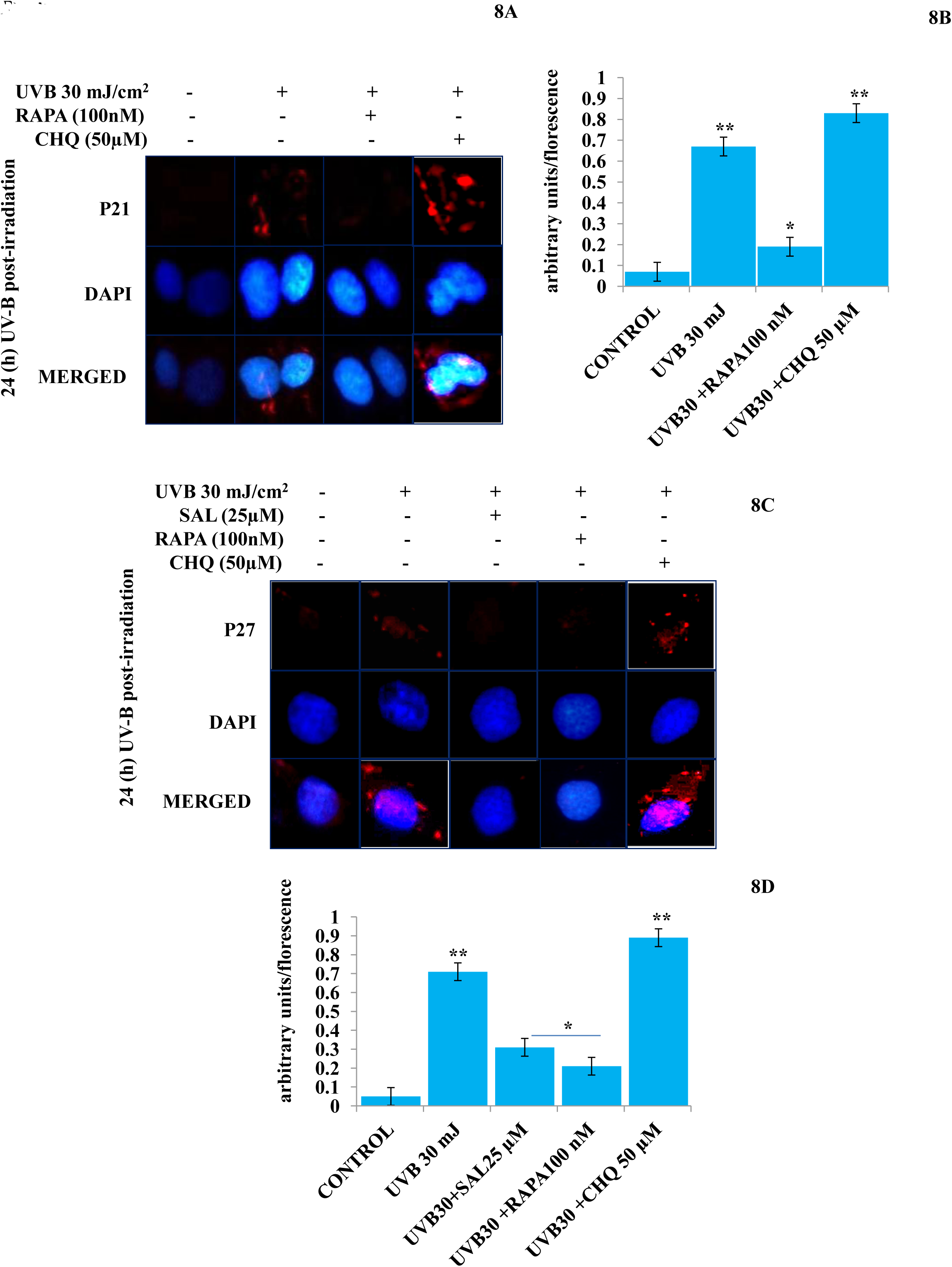
Rapamycin improves cell cycle regulation by regulating the expression of cell cycle regulatory proteins in immunoflorescence in UV-B exposed HDFs. (8A & 8B) Immunoflorescence analysis of P21 protein expression levels in UV-B 30 mJ/cm^2^ exposed HDFs in 24h UV-B post-irradiation in presence of Rapamycin (100nM) and Chloroquine (50µM). (8C & 8D) Immunoflorescence analysis of P27 protein expression levels in UV-B 30 mJ/cm^2^ exposed HDFs in 24h UV-B post-irradiation in presence of Salubrinal (25µM) Rapamycin (100nM) and Chloroquine (50µM). (*p≤ 0.05, **p≤ 0.01, were considered statistically significant).

### 3.9 ATG7 Silencing alleviates the DNA damage response in UV-B exposed HDFs

Autophagy related genes play an important role in the initiation and execution process of autophagy response. To check the specific role of particular autophagy related protein in UV-B induced photo-damage. We subject HDFs to ATG7 silencing in 6h UV-B post-irradiation. Western blotting analysis confirmed the silencing of ATG7 with 90% efficiency compared to control levels. UV-B 30mJ/cm^2^ exposure to HDFs increases the protein expression of ATG7 and BECN1 by 1 folds compared to control levels, (**p<0.01 for ATG7 in UVB 30mJ/cm^2^ exposed compared to control, **p<0.01 for BECN1 in UVB 30mJ/cm^2^ exposed compared to control). UV-B exposure to HDFs also induces the expression of key DNA damage response proteins pχH2AX and pP53 by 5 and 4.5 folds respectively, (#p<0.001 for pχH2AX in UVB 30mJ/cm^2^ exposed compared to control, **p<0.01 for pP53 in UVB 30mJ/cm^2^ exposed compared to control). ATG7 silencing in UV-B exposed HDFs significantly alleviates the DNA damage response as is evident from the decrease in the expression levels of DNA damage response proteins pχH2AX and Pp53 in western blotting by 2 folds, (**p<0.01 for pχH2AX in ATG7 ShRNA+UVB 30mJ/cm^2^ treated compared to UV-B exposed only, **p<0.01 for pP53 in ATG7 ShRNA+UVB 30mJ/cm^2^ treated compared to UV-B exposed only). ATG7 silenced only HDFs showed negligible effect on the modulation in expression level of autophagy and DNA damage marker proteins in western blotting analysis. Everolimus (200nM) treatment to ATG7 silenced cells upon UV-B exposure also decreased the expression levels of DNA damage response proteins to that of control levels, (*p<0.05 for pχH2AX in ATG7 ShRNA+UVB 30mJ/cm^2^+ EVER 200nM treated compared to UV-B exposed only, *p<0.05 for pP53 in ATG7 ShRNA+UVB 30mJ/cm^2^+ EVER 200nM treated compared to UV-B exposed only), (Fig. 9A & 9B). We confirmed our western blotting results through Immunoflorescence by looking out for pχH2AX foci and found that UV-B exposure to HDFs significantly induces the pχH2AX damage foci by 3 folds compared to control levels but are significantly decreased in ATG7 silenced UV-B exposed HDFs by 0.2 folds compared to that of UV-B only exposed, (**p<0.01 for pχH2AX in UVB 30mJ/cm^2^exposed compared to control, **p<0.01 for pχH2AX in ATG7 ShRNA+UVB 30mJ/cm^2^ treated compared to UV-B only exposed), (Fig. 9C & 9D).

**Fig. 9:**
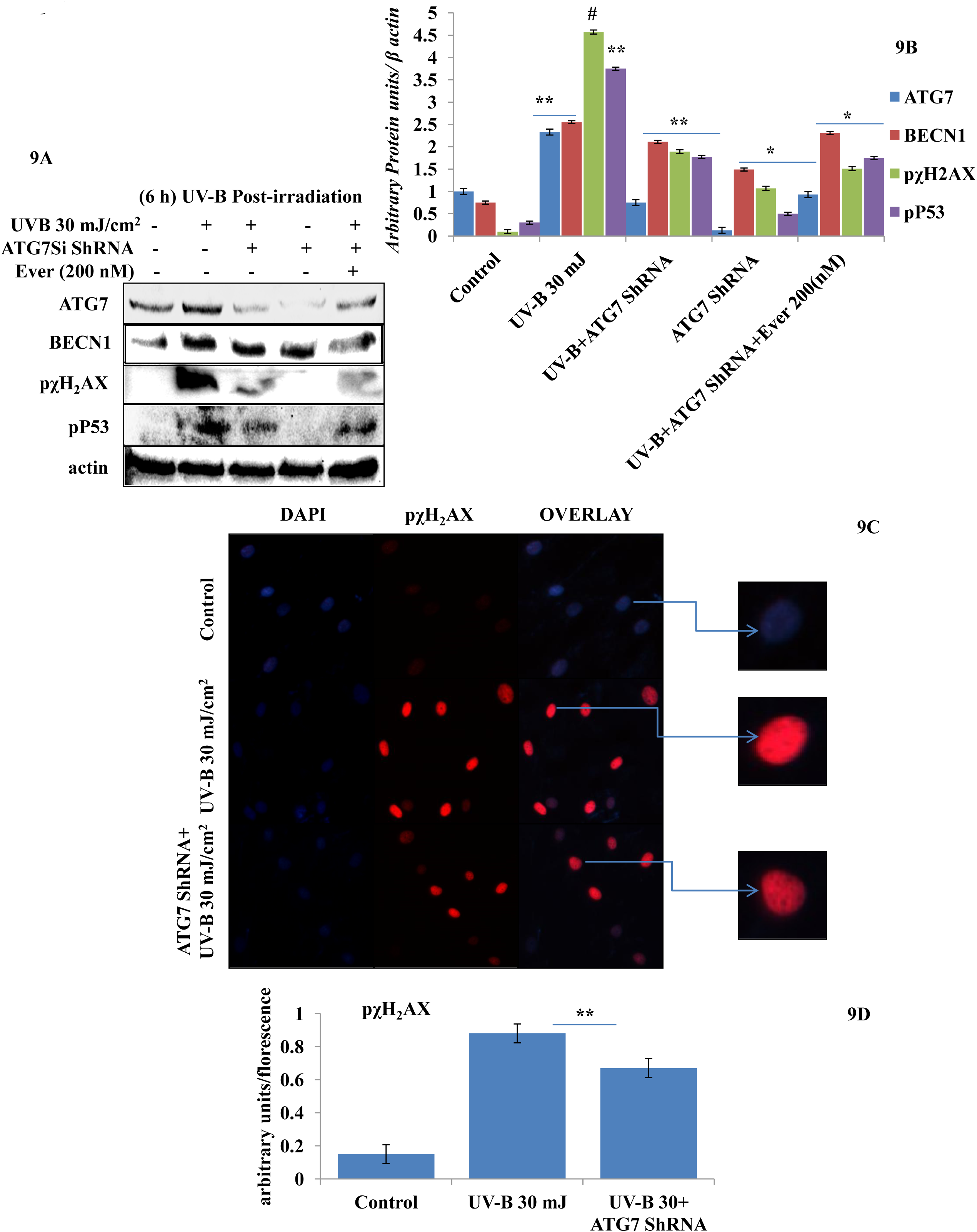
ATG7 Silencing alleviates the DNA damage response in UV-B exposed HDFs. (9A & 9B) Western blotting analysis of autophagy and DNA damage response proteins in ATG7 ShRNA treated HDFs in 6h UV-B post-irradiation in presence of Everolimus (200nM). (9C & 9D) Immunoflorescence analysis of DNA damage sensor protein pχH2AX in ATG7 ShRNA treated HDFs in 6h UV-B post-irradiation. (*p≤ 0.05, **p≤ 0.01, #p≤ 0.001 were considered statistically significant).

### 4. Discussion

Skin aging is a dynamic process and depends on both intrinsic factors such as genetics and hormones, as well as extrinsic factors like UV radiation and environmental pollutants [25]. UV radiation in particular is considered the most crucial factor for skin aging due to the process known as photoaging [26]. Incidences of skin cancer have increased in recent year’s world over, possibly due to increased exposure to solar ultraviolet radiation because of the depletion of ozone layer in stratosphere [27]. UV-B irradiation to skin stimulates diverse cellular and molecular responses like inflammation, reactive oxygen species (ROS) formation, Endoplasmic Reticulum (ER) stress response that ultimately lead to autophagy induction in skin cells aimed at restoring the internal homeostasis. Acute as well as chronic exposure of UV-B to skin lead to various perturbations that ultimately lead to aging-related signal transduction amplification resulting in skin damage and photoaging [28]. UV-B radiation is considered as the most mutagenic component of the UV spectrum reaching the earth’s surface and causes DNA damage in the form of cyclobutane pyrimidine dimers (CPD’s) and pyrimidine-(6-4)-pyrimidone photoproducts (6,4 PP’s) affecting DNA integrity, tissue homeostasis and also causes mutations in oncogenes and in tumour-suppressor genes [29]. Cells, have in defence, a natural, inbuilt and well established molecular response system known as DNA damage response (DDR) that checks any mutagenic insult to genome and repairs, immediately, to prevent tumor development and cancer progression. Any unrepaired part can lead to abnormal cell growth increasing the risk of cancer [30]. Autophagy is a cellular catabolic process that has roles in sensing nutrient stress during starvation conditions and cleanses cellular debris generated as metabolic byproduct [31]. Recent evidences from autophagy research in UV-B induced DNA damage conditions have provided novel insights and has been found to play a positive role in DNA damage recognition by nucleotide excision repair and controls p38 activation to promote cell survival under genotoxic stress [5, 32]. In another similar study, it has been found that Autophagic UVRAG promotes UV-induced photolesion repair by activation of the CRL4 (DDB2) E3 Ligase [33] citing positive role of autophagy in regulating UV-B induced DNA damage response in skin. We have recently reported that natural product based anti-oxidant molecule, Glycyrrhizic acid, alleviates oxidatively induced DNA damage response through improving autophagy levels in primary human dermal fibroblasts [19]. Despite these preliminary studies conducted so far in demystifying the role of autophagy in UV-B induced DNA damage response, the precise role of autophagy in regulating UV-B induced genotoxic stress is yet to be proved and warrants further studies to unearth the facts. In line with these findings, we planned the current study and hypothesized that autophagy might be playing a very crucial role in regulating UV-B –induced DNA damage response. ELISA based results reveal that DNA photo-adducts (CPD’s and 6, 4 PP’s) are the immediate byproducts of oxidative damage in UV-B exposed HDFs in 6h UV-B post-irradiation. Improving autophagy flux with Rapamycin (100nM) and on relieving UV-B – induced ER stress response with Salubrinal (25µM), significantly reduces the formation of both CPD’s and 6, 4 PP’s. Autophagy inhibitor Chloroquine (25 µM) treatment to HDFs has little effect on the formation of CPD’s whereas, it increases the formation of 6, 4 PP’s in UV-B treated HDFs (Fig. 1A), citing that autophagy induction positively regulates the formation of DNA photo-adducts in UV-B induced photo-damage. These results were confirmed through silencing autophagy cargo protein P62 to confirm whether improving autophagy levels also improves the UV-B -induced photo-damage response in HDFs. We found that it significantly impacts the formation of photo-adducts moreso CPD’s than that of 6, 4 PP’s and increases the formation of CPD’s and 6, 4 PP’s whereas P62 silenced only cells have same effect as that of control indicating that autophagy plays very critical role in UV-B induced photo-damage (Fig. 2B). Similar effects were obtained in western blotting as well as in immunoflorescence on checking the effect of improving cellular autophagy levels on CPD’s formation, further revealing that autophagy regulates DNA damage in UV-B exposure to HDFs, (Fig. 1D & 1E, Fig, 1F & 1G). Our results further reveal that Rapamycin (100nM) treatment to HDFs has no significant effect on restoring the cellular viability in UV-B 30mJ/cm^2^ exposed HDFs whereas Chloroquine 50µM treatment significantly reduces the cell viability from 65% in UV-B 30mJ/cm^2^ only to 60% in UV-B30+CHQ (50µM) treated, indicating that on inhibiting autophagy potentiates the cell death potential of UV-B exposure to HDFs (Fig. 1C). It is a given fact that UV-induced skin damage triggers a cascade of response signaling pathways, including cell cycle arrest, DNA repair, and, if left unrepaired, can lead to apoptotic events [34]. Oxidatively induced DNA damage response is the hallmark of UV-B –induced skin photo damage that ultimately leads to genotoxic stress response [35]. We found that UV-B treatment to HDFs induces TUNNEL positive cells in an intensity dependent manner in 6h UV-B post-irradiation. Rapamycin treatment (100nM) significantly reduces the florescence of TUNNEL positive cells in UV-B 30mJ/cm^2^ exposed HDFs whereas Chloroquine treatment (50µM) to HDFs increases the florescence of TUNNEL positive cells compared to UV-B only exposed, (Fig. 2A & 2B). AO/EtBr co-staining also reveals increase in the florescence of apoptotic nuclei in microscopic studies in UV-B exposed HDFs indicating nuclear damage on UV-B exposure. Salubrinal (25µM) and Rapamycin (100nM) treatment significantly quenched the florescence of apoptotic nuclei to that of control levels whereas Chloroquine (50µM) treatment increases the florescence in EtBr co-staining compared to UV-B levels. These results clearly indicate that Oxidative stress, ER stress and autophagy process are intricately interconnected and that autophagy has a very critical role in alleviating the stress response in cells, (Fig. 2C & 2D). Recent works have revealed that genotoxic stress is a trigger for autophagy and autophagy regulates repair of UV-induced DNA damage. It was found previously that knockdown of autophagy genes such as AMPK, Atg5, Atg7, Atg12, and Atg14 impairs the repair of UVB-induced DNA damage [36]. Our results reveal that autophagy response is the immediate molecular event following UV-B exposure to HDFs and is a time dependent response induced in an intensity dependent manner 1 to 6h post-UV-B –irradiation, after then fades as evident from the western blotting analysis of key autophagy marker proteins but depicts impaired autophagy flux, (Fig. 3A, 3B, 3C, 3D & 3E). GFP-RFP-LC3B puncta assay which combines the ability to monitor the various stages of autophagy (through LC3B protein localization) also reveals that UV-B exposure to HDFs induces impaired flux as evident from the lower GFP to RFP conversion, but Rapamycin treatment significantly rescues the UV-B exposed HDFs from photo-damage effects through improving autophagy levels as is clear from improvement in conversion ratio of GFP to RFP depicting significant PUNCTA positive cells. Further Chloroquine treatment to UV-B 30mJ/cm^2^ exposed HDFs shows least GFP-RFP-LC3B PUNCTA positive cells and has also stalled the conversion of GFP to RFP that depicts progression of autophagy and autophagosome maturation (Fig. 3F & 3G). Immunoflorescence results also reveal that autophagy induction is an immediate event in 1h UV-B post-exposure to HDFs as evident from the western blotting results of P62 and LC3B. Considering that autophagy and DNA damage response execute parallel in UV-B photo-response in skin, we got similar results for DNA damage response proteins pχH2AX and pP53 in Immunoflorescence showing that when there is intense cell damage by UV-B exposure, autophagy is the immediate event to retaliate and repair the photo-products, (Fig. 3H & 3I). Moreover, oxidative stress induced by the formation of ROS upon UV-B exposure to skin also induces autophagy and primary ROS is the main oxidative damage causing agent in UV-B exposed cells [37]. We found that treatment with Salubrinal as ER stress reliever and Everolimus as autophagy inducer significantly alleviates the ROS levels to half produced in response to UV-B exposure to skin cells indicating that improving autophagy levels has a descent role in regulating oxidative stress response and that ER stress and oxidative stress are mutually related to each other disturbing the cellular homeostasis (Fig. 3J & 3K). Intense UV-B exposure to HDFs induces genomic damage and in response cell cycle arrest is critical at providing ample time gap for DNA damage recognition and subsequent execution of repair process. UV-induced DNA damage activates the sensors ataxia telangiectasia mutated (ATM) and ataxia telangiectasia and Rad3-related (ATR) to trigger cell cycle arrest via p53 stabilization [38] and also phosphorylates checkpoint kinase 1 (Chk1) to activate checkpoints at the G1, S, and G2/M phases. Damage related protein DDB2 have been shown to facilitate the recruitment of ATM and ATR to sites of DNA damage and promote the activation of cell cycle arrest pathways [39–41]. We found that UV-B induced DNA damage response is both a time and intensity dependent event as is evident from the western blotting analysis of DNA damage response proteins (Fig. 4A to 4D). The expression level of pAKT, which has crucial role in pro-survival signaling and also inhibits apoptosis in UV-B response is also significantly upregulated in UV-B 30mJ/cm^2^ exposed HDFs (Fig. 4A & 4E). pχH2AX foci are immediately induced upon DNA damage, sensing damage and facilitating repair process. Here we found that UV-B 30mJ/cm^2^ exposure to HDFs induces the expression of pχH2AX foci in Immunoflorescence. Rapamycin (100nM) treatment to UV-B exposed HDFs rescues HDFs from DNA damage as is evident from the decreased expression of pχH2AX whereas Chloroquine (50µM) and Bafilomycin A1 (100nM) treatment show enhanced expression levels of pχH2AX nuclei in UV-B 30mJ/cm^2^ exposed HDFs indicating that autophagy positively regulates the damage responsive wing in UV-B exposed HDFs, (Fig. 4F & 4G). Immunoflorescence of DDB2 also reveals upregulated levels of expression in UV-B 30mJ/cm^2^ exposed HDFs. Here also, Rapamycin (100nM) significantly brought the DDB2 level to that of control whereas Chloroquine (50µM) treatment drastically increases the expression level of DDB2 protein further, than in UV-B exposed HDFs, (Fig. 4H & 4I) which indicates that DNA damage response dwindles if autophagy levels are improved in UV-B exposed HDFs and the pro-survival capacity of cells is enhanced due to clearing of damage incurred due to UV-B exposure to HDFs.

Previously, we have reported that oxidative stress mediated Ca^2+^ release manifests endoplasmic reticulum stress leading to unfolded protein response in UV-B irradiated human skin cells [18]. In another study, it was reported that Salubrinal protects human skin fibroblasts against UVB-induced cell death by blocking endoplasmic reticulum (ER) stress and regulating calcium homeostasis [42]. Here, we found that Salubrinal (10-30µM), an eif_2_α inhibitor, significantly restores the cellular viability in MTT assay in UV-B 10-30mJ/cm^2^ exposed HDFs (Fig 5A) and was found to be safe upon 30 µM. Salubrinal (25µM) significantly reduces ER stress response (5G & 5K) and also alleviates the DNA damage in 6h post-UV-B 30mJ/cm^2^ exposed HDFs as evident from western blotting analysis of DNA damage marker proteins (5H & 5L) but not significantly in 24h UV-B post-irradiation to HDFs at 20µM working concentration, (Fig 5B to 5F). ER calcium leakage is the immediate post-event following UV-B exposure to HDFs as reported earlier [18]. We found that UV-B mediated ER calcium leakage is significantly prevented by Salubrinal (25µM) and Rapamycin (100nM) treatment to UV-B exposed HDFs in 6h UV-B post-irradiation, whereas Chloroquine (50 µM) and Bafilomycin A1 (100nM) failed to prevent the calcium depletion from ER in microscopic analysis (Fig. 5I & 5J).

Previous studies have reported that p62 modulates the intrinsic signaling of UVB-induced apoptosis [43] and that autophagy performs crucial role in promoting cell survival under genototoxic stress and helps in preventing tumorigenesis. Degradation of P62 has been found to mediate an important tumor suppressive function of autophagy and autophagy-deficient conditions have shown enhanced P62 accumulation and acts as a signaling hub by forming interactions with a number of pro-tumorigenic proteins, thereby promoting tumorigenesis [44]. Here we found that autophagy blockage via P62 silencing shows enhanced DNA damage in 6h UV-B 30mJ/cm^2^ post-irradiation as is clear in western blotting analysis of DNA damage response proteins, (Fig. 6A & 6B) and similar results were obtained in immunoflorescence of pχH2AX foci in confocal microscopy which significantly increase in P62 silenced cells. Rapamycin (100nM) dwindles the expression of damage sensor pχH2AX to that of control levels whereas Chloroquine (50µM) significantly increases the expression of pχH2AX in immunoflorescence indicating that autophagy induction regulates the UV-B induced damage and imparts pro-survival capability to UV-B exposed HDFs, thereby reducing chances of cancer development, (Fig. 6C & 6D). Earlier, it has been reported that PTEN which otherwise activates autophagy is inhibited by Sestrin 287 and in response to UV-B exposure impairs the GG-NER by downregulating XPC transcription. However, it is still elusive to conclude that whether downregulation of autophagy has a significant role in PTEN-regulated DNA damage repair in response to UV-B exposure to skin cells [45–47]. PTEN/AKT pathway is the crucial tumor suppressor pathway that promotes cell survival under genotoxic stress and earlier studies have found that ERK/AKT-dependent PTEN suppression promotes survival of epidermal keratinocytes under UV-B exposure. Our results are in complete agreement with these initial findings as P62 silencing disturbs the PTEN/pAKT pathway in UV-B exposed HDFs in 6h UV-B post-irradiation by decreasing the PTEN protein expression level but increases the pAKT levels compared to UV-B, (7A & 7B). Similar results were obtained in immunoflorescence of PTEN and pAKT, (Fig. 7C & 7D, 7E & 7F, respectively) which indicate that autophagy also regulates tumor suppressor activity under UV-B induced genotoxic stress. Further, Rapamycin (100nM) improves the PTEN/pAKT mediated tumor suppressor pathway whereas Chloroquine (50µM) potentiates the UV-B response to HDFs in microscopy backing our silencing results.

The massive accumulation of DNA lesions within the cells under different genotoxic stimuli interferes with replication process, prompting cells to stop division and to repair damaged DNA [48]. Cell cycle regulator proteins this way play a very important part in quality control of cells and to sense any external insult and prevent mutations. We found that Salubrinal (25µM) and Rapamycin (100nM) improves the fate of cell cycle regulator proteins P21 (Fig 8A & 8B) and P27 (Fig 8C & 8D) in immunoflorescence in 24h UV-B post-irradiation through regulating autophagy response whereas Chloroquine (50 µM) further worsens the damage regulation by increasing the expression of P21 and P27 which indicates that autophagy also plays its part in regulation of cell cycle proteins under genotoxic stress conditions.

Earlier studies have revealed unexpected consequence of Atg7 gene deletion in the suppression of UVB-induced inflammation and tumorigenesis through the PTGS2-PGE2 pathway and epidermis-specific deletion of Atg7 has been found to protect against UVB-induced sunburn, vascular permeability, and skin tumorigenesis. Moreover, ATG7 deletion has been found to regulate PTGS2 expression and UVB-induced skin tumorigenesis through regulating the AMPK and ER pathways, [49]. Our results are in complete agreement with these previous findings because ATG7 silencing in UV-B exposed HDFs alleviates the DNA damage induced in 6h UV-B post-irradiation to HDFs as is evident from the western blotting analysis of key DNA damage marker proteins. Everolimus (200nM) also rescues the HDFs from UV-B induced DNA damage under ATG7 silencing conditions, (9A & 9B). Similar results were obtained in immunoflorescence by looking for pχH2AX expression levels which are significantly reduced in ATG7 silenced HDFs compared to UV-B exposed only, (Fig. 9C & 9D) demonstrating an unexpected consequence of Atg7 silencing and its role in the suppression of UVB-induced DNA damage response, but warrants more studies to clearly understand the complexity and intricacies involved in demystifying the role of autophagy particularly of ATG7 gene deletion in regulation of UV-B induced genotoxic stress response.

## 5. Conclusion

Above findings provide some critical insights that indicate the regulatory and functional role of autophagy in regulation of UV-B mediated skin photo-damage. These findings further reveal that cellular autophagy levels are critical in sensing and repairing UV-B –induced DNA damage and is crucially involved in DNA damage response under UV-B induced photo-damage conditions. Oxidative stress-ER stress and DNA damage response mechanisms are tightly regulated and that autophagy is involved in regulation of all the three pathways and promotes pro-survival capacity of cells under genotoxic stress conditions notably under UV-B –induced DNA damage. Moreover, our findings support the potential role of autophagy pathway to be explored as promising therapeutic and cosmeceutical target in the protection of skin cells from UV-B mediated photodamage disorders but warrants further studies to demystify the molecular association between autophagy and DNA damage response in UV-B –induced skin photo-damage.

## Acknowledgements

Senior Research Fellowship (SRF) to author SAU by Department of Science and Technology (DST), Government of India Vide No. IF-160982 is acknowledged. Authors are also thankful to Director CSIR-Indian Institute of Integrative Medicine, Jammu for funding this work vide Project No. MLP-1003 and Department of Biotechnology (DBT), Ministry of Science and Technology, Government of India, New Delhi, India vide project No. GAP-2166. Authors also thank all the members of Experimental Toxicology Laboratory, PK-PD & Toxicology Division for their logistic support and help in carrying out this study.

## Author contributions

SAU performed the experiments. SAT conceived and developed the hypothesis, supervised the research work and arranged the research funding for the work. SAU and SAT planned the experiments, analyzed the data and wrote the manuscript. All the authors provided critical feedback and helped shape the research and analysis of data.

## Abbreviations

UV-B: Ultraviolet-B
HDFs: Human Dermal Fibroblasts
DNA: De-oxy ribo Nucleic acid
DDR: DNA Damage Response
ROS: Reactive Oxygen Species
ER: Endoplasmic Reticulum
CPD: Cyclobutane Pyrimidine Dimers
6, 4 PP: Pyrimidine-(6-4)-pyrimidone
NER: Nucleotide Excision Repair
BER: Base Excision Repair
MR: Mismatch Repair
NHEJ: Non homologous end joining
GA: Glycyrrhizic acid
MTT: 3-(4, 5-dimetylthiazol-yl)-diphenyl tetrazolium bromide
H2DCFDA: Dichlorofluorescin diacetate
RAP: Rapamycin
SAL: Salubrinal
CHQ: Chloroquine
BAF A1: Bafilomycin A1
AO: Acridine Orange
EtBr: Ethidium Bromide
DPBS: Dulbecco’s phosphate buffered saline
FBS: Fetal bovine serum
DMSO: Dimethyl Sulfoxide
GFP: Green florescent protein
RFP: Red florescent protein
ELISA: Enzyme-linked immunosorbent assay
ICC: Immuno cytochemistry

